# Alkyladenine DNA glycosylase regulates TDG-mediated demethylation and neuronal gene expression by competitive substrate binding

**DOI:** 10.64898/2026.11.15.766739

**Authors:** Ezgi Merdane Aksu, Kayla Grooms, Andreas Gjøsæther, Nicholas Ertresvaag Zachariadis, Arnar Flatberg, Matthew A. Schaich, Rabina Dumaru, Cathrine Broberg Vågbø, Diana L. Bordin, Pål Sætrom, Bennett Van Houten, Barbara van Loon

## Abstract

Endogenous single-strand DNA breaks (SSBs) that arise during active DNA demethylation, regulate neurodevelopmental gene expression. Thymine (TDG) and alkyladenine (AAG) DNA glycosylases recognize oxidized methylcytosines. While TDG is required for excision, AAG’s role in active demethylation and neurodevelopment remains unknown. Using mouse neurospheres, hippocampus, and human induced neurons (iNs), we show that AAG is needed for neurogenesis. Loss of AAG contributes to delayed neurodevelopmental and premature synaptic gene expression, evident from single-cell RNA sequencing of differentiating iNs. Synaptic gene expression is co-regulated by both AAG and TDG, with loss of single glycosylase inducing an opposing effect, implying competition. Single-molecule and biochemical assays directly demonstrate this competition, with AAG inhibiting TDG on 5fC. Without AAG, TDG has higher affinity, resulting in reduced global 5fC and increased SSB intermediates. In summary, AAG substrate competition plays an important role in fine tuning active demethylation, thereby regulating gene expression and DNA breaks during neurogenesis.

## Introduction

Neurons are key units for cognition and behavior. Neuronal activity has essential functions in formation and refinement of neuronal circuits, and is of pivotal importance for learning and memory^1^. In line with this, the neuronal activity continuously regulates synaptic transmission and drives synapse formation through activation of distinct gene expression profiles^2, 3^. Disruption of neuronal activity-dependent gene regulation leads to a wide range of neuropathological conditions^4^.

Interestingly, the neuronal activity-triggered gene expression changes are largely dependent on induction of DNA breaks in the gene promoters^5^. Despite being mostly considered harmful threats to genome integrity, DNA breaks and the associated DNA repair pathways are emerging as important regulators of neuronal functions. For instance, levels of endogenous DNA breaks are increased at neuron-specific genes of neuronal stem and progenitor cells^6–8^. The increase is required for proper neuronal gene expression, and subsequently for neuronal differentiation. In line with this, the single strand DNA breaks (SSBs) were shown to contribute to neuronal fate commitment^9–11^. Recently, base excision repair (BER) of oxidized methylcytosines, occurring during active demethylation, has been identified as a primary source of these SSBs^9^.

The oxidized methylcytosine, like 5-hydroxymethylcytosine (5hmC), is highly enriched in the nervous system and is an epigenetic mark that modulates gene expression^12, 13^. Other types of oxidized methylcytosine arise during active DNA demethylation, which is especially pronounced in the developing brain^14^. During active DNA demethylation, 5-methylcytosine (5mC) is oxidised by the ten–eleven translocation (TET) family of methylcytosine dioxygenases (TET1, TET2, and TET3) to 5hmC^15^, which can be further oxidised to 5- formylcytosine (5fC) and 5-carboxylcytosine (5caC). The active DNA demethylation cycle is completed by 5fC and 5caC removal through the base excision repair (BER) pathway^16–18^. BER is initiated by DNA glycosylases that recognize and excise chemically modified substrate bases, thus generating abasic (AP) sites, which are subsequently processed by apurinic/apyrimidinic endonuclease 1 (APE1)^19^. The resulting single-strand break intermediates are sensed by poly-ADP-ribose polymerase 1 (PARP1) and XRCC1 scaffold protein^20^, which coordinate the recruitment of DNA polymerase β and DNA ligase III for gap filling and ligation, respectively, consequently the completion of short patch BER.

In mammals, BER is initiated by one of eleven DNA glycosylases with broad substrate specificities^21^. Thymine DNA glycosylase (TDG) has central functions in active demethylation and is a primary enzyme for 5fC and 5caC binding and removal^22, 23^. Besides TDG, alkyladenine DNA glycosylase (AAG, aka MPG) is the only other DNA glycosylase that directly binds 5fC^24, 25^. While substrate overlap among DNA glycosylases is widely reported, we still lack a clear understanding of its biological consequences. The functional importance of AAG binding to 5fC furthermore remains unclear, since this DNA glycosylase cannot remove this base.

Canonically, AAG functions as key glycosylase for removal of methylated base lesions, such as 7-methylguanine (7meG) and 3-methyladenine (3meA). Increasing evidence suggests an important role for AAG beyond canonical genome maintenance. For example, we showed that AAG directly associates with active transcription, impacting the expression of neuronal genes^26^, and consistent with its interactions with several transcription factors, including methyl binding domain (MBD1) and estrogen receptor (ER)^27, 28^. In addition to 5fC, AAG has been identified as a “reader” of 5hmC^24^. Our recent findings indicate that AAG status directly impacts 5hmC and DNA break levels in mouse hippocampus, altering neuronal transcriptome, with direct consequences for brain functions and behavior^29^. Despite the emerging roles in neuronal physiology, the functional role of AAG in active demethylation and neurodevelopmental gene regulation remains unknown.

In this study, we show that AAG promotes neurogenesis, with its loss manifesting in reduced neuronal markers in both *Aag^-/-^* mouse neurospheres and hippocampus, and human induced neurons (iNs). During neuronal differentiation lack of AAG results in downregulation of neurodevelopmental and upregulation of synaptic genes, as evident from the single-cell RNA sequencing across iNs developmental timeline. Interestingly, lack of TDG in the same model system has an opposing effect, resulting in downregulation of synaptic genes, thus suggesting negative cross-regulation. Consistent with this observation, assessment of DNA demethylation intermediates in AAG^-/-^ cells uncovered significantly reduced global genomic 5fC and 5caC levels, 15% and 30% lower, respectively. Thus, suggesting that absence of AAG promotes TDG activity on shared substrates such as 5fC. This is further evident from the single-molecule and biochemical studies, which for the first time visualize competition between two DNA glycosylases, revealing notably higher TDG affinity for 5fC in the absence of AAG. In summary, our findings provide evidence for a functional substrate competition between DNA glycosylases that regulates active DNA demethylation and consequently gene expression programs, important to ensure unperturbed neurodevelopment.

## Results

### Neurogenesis is impaired in AAG lacking mouse neurospheres and human induced neurons and is accompanied by reduced level of DNA breaks

Processing of modified DNA bases by AAG, and DNA glycosylases in general, is emerging as one of the essential mechanisms for regulation of neural gene expression, relevant for unperturbed brain functions and behavior^9, 29–33^. However, it remains, largely unclear whether AAG primarily exerts its functions during neurodevelopment or in mature neurons. To address this and to assess the impact of AAG status on neural progenitor cells (NPCs), a mouse brain neurosphere model system was used. Neurospheres were obtained upon propagation of NPCs isolated from brains of wild type (WT) and Aag knock-out (*Aag^-/-^*) pups at birth. Both the NPCs self-renewal and proliferation capacities remained unaltered upon Aag loss and were comparable between *Aag^-/-^* and WT NPCs (Supplementary Fig. 1a, b). Next, to investigate the impact of Aag on NPC differentiation potential, spontaneous NPC differentiation towards neurons and glia was initiated, followed by immunofluorescence (IF) analysis. Neurons and glia were identified by using class III β-tubulin (Tuj1), a marker of immature postmitotic and differentiated neurons, and glial fibrillary acidic protein (Gfap), an intermediate filament highly expressed in astrocytes. Interestingly, the signal of Tuj1 positive neuronal cells was significantly reduced upon differentiation of *Aag^-/-^* NPCs, when compared to WT cells (Fig. 1a, b). In contrast the expression of glial marker Gfap was comparable between WT and *Aag^-/-^* cells (Fig. 1a, c). To further investigate Aag effects on the whole tissue level, IF analysis targeting mature neuron-specific microtubule-associated protein 2 (Map2) and Gfap in the hippocampus of young mice at age postnatal day 5 (P5) was performed (Supplementary Fig. 1c). Mature neuronal marker Map2 was significantly reduced in the *Aag^-/-^* compared to WT hippocampus (Supplementary Fig. 1d). Notably, Map2 protein signal was significantly decreased in hippocampal regions CA1 and CA3 (Supplementary Fig. 1f, g), which are associated with glutamatergic neuron populations, and not in the dentate gyrus (DG) (Supplementary Fig. 1e). Similar to the findings in the neurospheres, Gfap signal was not affected by Aag loss in P5 mouse hippocampus (Supplementary Fig. 1h-k). These results thus suggest that lack of Aag in mice impairs neuronal, but not glial differentiation.

**Figure 1.**
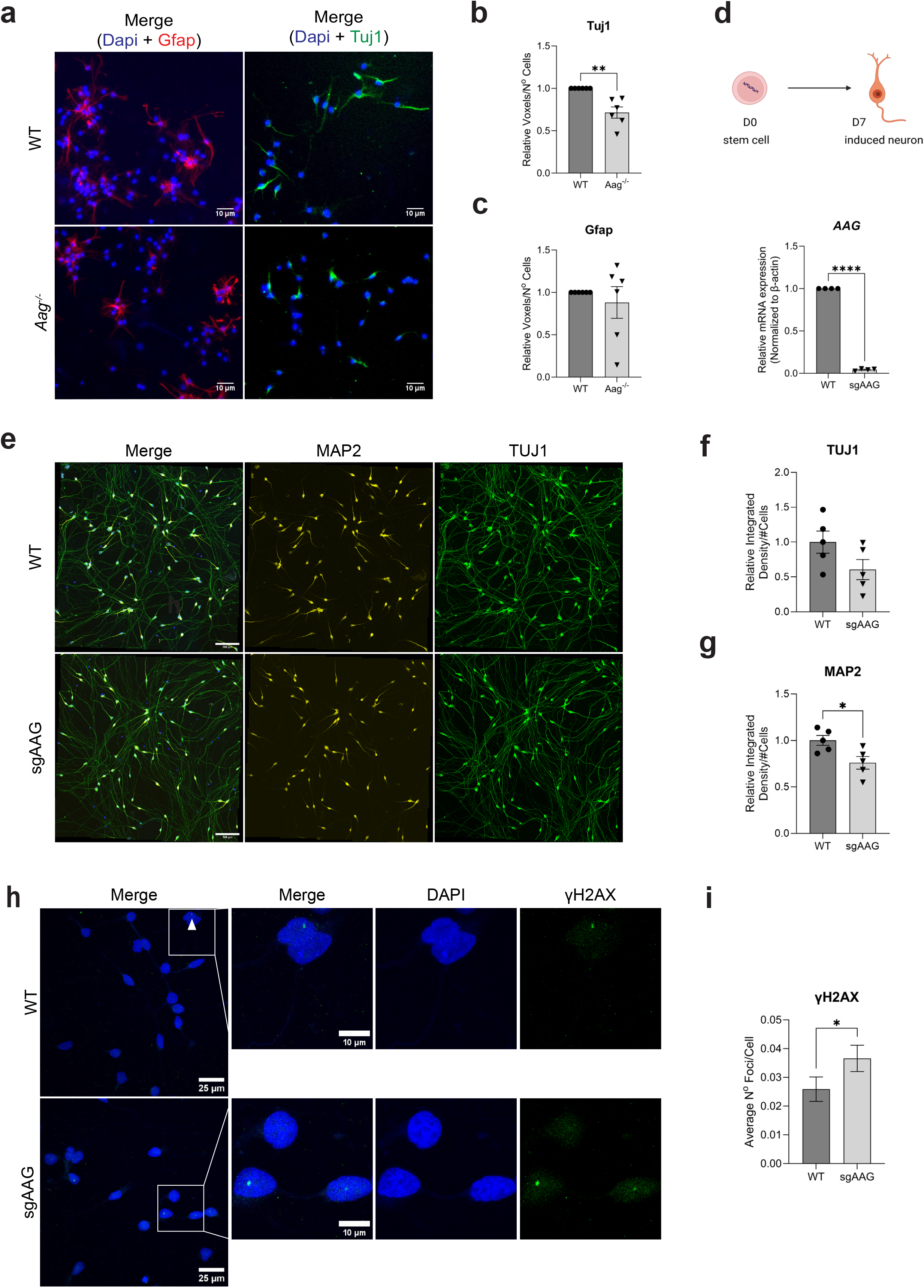
Loss of AAG leads to impaired neurogenesis in mouse neurospheres and human induced neurons, accompanied by reduced DNA break levels. **a** Representative immunofluorescence images of differentiated neuronal progenitor cells (NPCs) from wild type (WT) and Aag knock-out (*Aag^-/-^*) neurospheres stained for the neuronal marker Tuj1, glial marker Gfap and DAPI. (Scale bar = 10 µm). **b-c** Quantification of Tuj1 (b) and Gfap (c) signals from four independent experiments as the one depicted in (a). Quantification represents the number of voxels from the fluorescent signal normalized by the number of DAPI positive cells (n = 4). **d** Relative AAG mRNA levels in non-targeting sgRNA (WT) and AAG targeting sgRNA (sgAAG) induced neurons (iNs) at D7 of differentiation, normalized to β-actin mRNA. **e** Representative immunofluorescence images of MAP2 and TUJ1 staining in non-targeting sgRNA (WT) and AAG targeting sgRNA (sgAAG) induced neurons (iNs) at day 7 (D7) of differentiation. Scale bar 100 µm. **f-g** Quantification of relative TUJ1 (f) and MAP2 (g) signal intensities in D7 iNs (n= 4-5 coverslips, from 3 independent differentiations). **h** Representative immunofluorescence images of the γH2AX signal in the D7 WT and sgAAG iNs (n = 1545 and 1831 cells, respectively; scale bar = 25 µm; scale bars magnified images = 10 µm). **i** Quantification of three independent experiments as the one in (h). All data represented as mean ± SEM, *p ≤ 0.05, **p ≤ 0.01, ****p ≤ 0.0001, two-tailed Student’s t-test. Cell illustrations created in BioRender by Aksu E.M. https://BioRender.com/xhsb61e

To determine the relevance of these findings for human development, human cortical neurons were generated next by using a human induced pluripotent stem cell (hiPSCs) line carrying dCas9 transcriptional repression system and an inducible *NGN2* promoter^34, 35^. Using lentiviruses encoding AAG-targeting sgRNA (sgAAG) we first silenced AAG expression; a non-targeting sgRNA (WT) served as a control (Supplementary Fig. 2a, b). Neuronal induction was initiated in WT and sgAAG hiPSCs with doxycycline for 3 days (D3), the differentiating neurons collected on D7 and subjected to IF analysis of neuronal markers (Fig 1d, e). In line with the findings obtained in mouse neurospheres and hippocampus, levels of both immature TUJ1 and mature MAP2 neuronal proteins were reduced up to two-fold in sgAAG compared to WT iNs (Fig 1f, g). It has been reported that generation of DNA breaks through activity of repair proteins influences cell fate decisions and promotes neuronal differentiation^9^. To determine if loss of AAG affects levels of DNA breaks in iNs, we next analyzed number of phosphorylated histone variant H2AX (γH2AX) foci within the nuclei of iNs. Surprisingly, sgAAG iNs had significantly more γH2AX foci compared to WT neurons (Fig. 1h, i). In summary, these findings propose that AAG loss impairs neurogenesis across species and contributes to increased number of DNA breaks in neurons.

### Loss of AAG alters temporal gene expression during neuronal differentiation

Timely activation of transcriptional programs through induction of DNA breaks is central for efficient neuronal differentiation. To identify the impact of AAG on temporal dynamics of gene expression in differentiating neurons, WT and sgAAG induced neurons were collected across different maturity stages (D4, D7, and D14) and subjected to single cell RNA sequencing (scRNA-seq) (Fig. 2a). Differences in the maturity of iNs from D4-14 were assessed by RT- qPCR and compared to stem cell state at D0. As expected, expression of stemness marker OCT4 was only present at D0, whereas levels of PAX6, an NPC marker that drives the generation of excitatory neurons, peaked at D7 (Supplementary Fig. 2c, d). Early neuronal marker DCX and mature neuronal marker MAP2, as anticipated, increased over time with highest levels observed at D14 (Supplementary Fig. 2e, f). Subsequent scRNA-seq analysis of D4, D7 and D14 iNs identified 4790 genes with altered expression trajectories in sgAAG cells when compared to WT iNs (Fig. 2a). Subsequent clustering of the temporal expression patterns revealed two major gene clusters: Cluster 1 containing 2683 genes with preterm activation in sgAAG iNs (Fig. 2b) and Cluster 2 with 236 genes whose expression was delayed in sgAAG iNs (Fig. 2c). Interestingly, the majority of upregulated Cluster 1 genes were associated with synaptic vesicle endocytosis and protein regulation processes (Fig. 2d), as determined by DAVID gene ontology (GO) enrichment analysis^36^. Meanwhile the downregulated Cluster 2 genes were primarily enriched in nervous system development and neuronal differentiation processes (Fig. 2e). Subsequent pseudotime and RT-qPCR analyses of the most significantly altered genes associated with synaptic vesicle endocytosis and neuronal differentiation and development processes, *RIMS4* (Fig. 2f, h) and *RBFOX3* (Fig. 2g, i) respectively, further confirmed preterm activation and repression, thus mirroring Cluster 1 and 2 expression patterns (Fig. 2b, c). That loss of AAG hampers neuronal differentiation genes was further supported by *MAP2* and *DCX* RT-qPCR analysis, which revealed significantly reduced DCX and MAP2 expression in sgAAG, compared to WT iNs (Supplementary Fig. 2e, f). Taken together, these results strongly suggest that the absence of AAG alters temporal neuronal gene expression programs, leading to preterm activation of synaptic transmission and repression of neurodevelopmental and differentiation genes.

**Figure 2.**
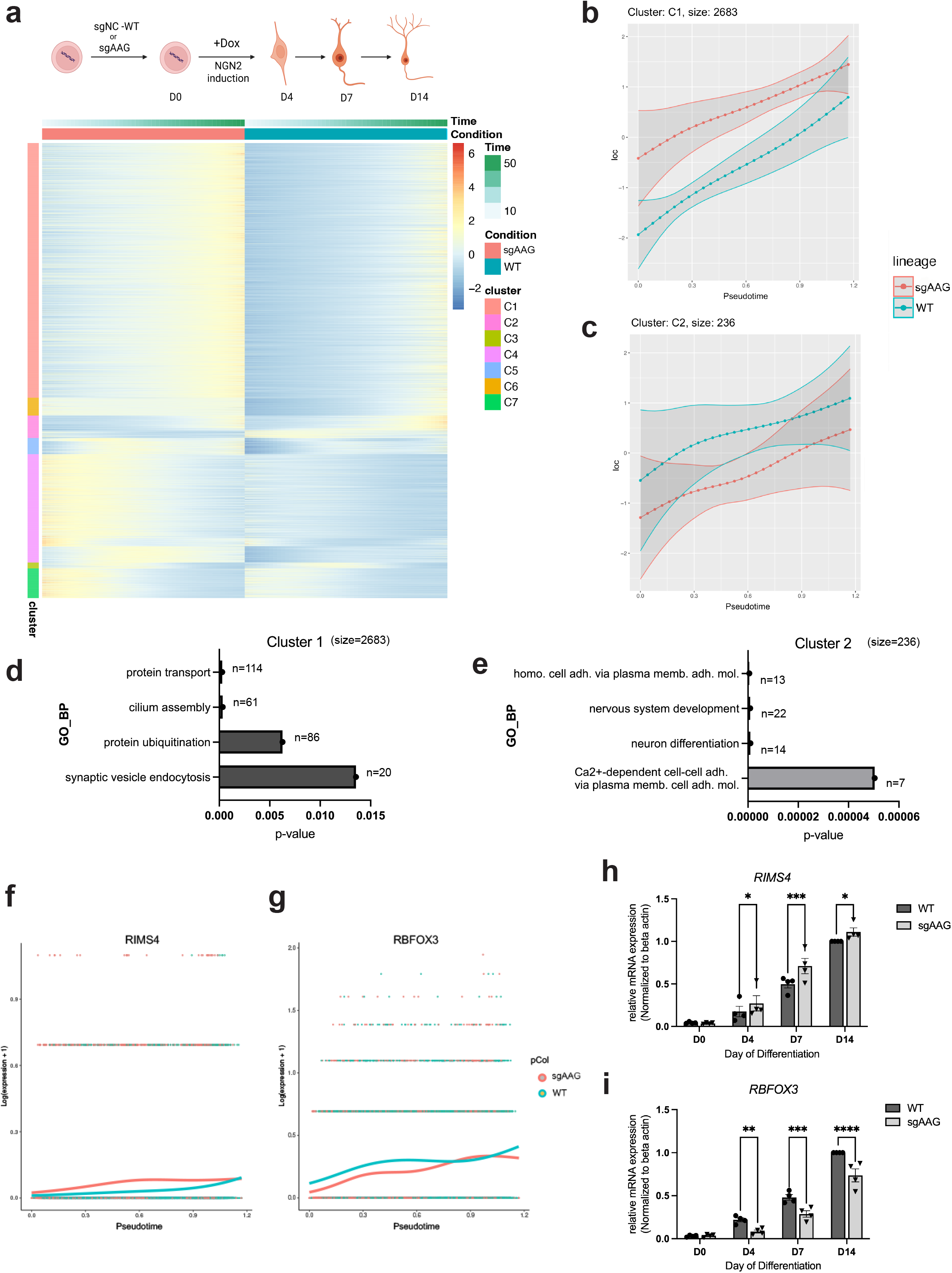
Neuronal gene expression is dysregulated in AAG lacking human induced neurons (iN). **a** Heat map of differential gene expression along differentiation pseudotime in non-targeting sgRNA (WT) and AAG targeting sgRNA (sgAAG) iNs; day (D) 4 to D14 of differentiation. Color scale represents z-score of gene abundance. Genes with significant pseudotime-dependent expression are clustered using the degPatterns into co-expression modules, clusters (C1-7). **b-c** Differential expression along pseudotime of genes in WT and sgAAG iNs from cluster C1 (b) and cluster C2 (c); revealing gene modules activated prematurely C1, or late C2 upon AAG loss in iNs. **d-e** Top four most significant biological processes (BP) gene ontology (GO) terms, determined by the Database for Annotation, Visualization and Integrated Discovery (DAVID), for C1 (d) and C2 genes (e). **f-g** Example of fitted expression profiles for synaptic *RIMS4* (f) and neuronal *RBFOX3* (g) genes, from C1 and C2 respectively, along the WT and sgAAG iN differentiation timeline. **h-i** RT-qPCR gene expression analysis confirming altered *RIMS4* (h) and *RBFOX3* (i) expression levels along WT and sgAAG differentiation, from D0 stem cells towards iNs (D4-D14). All error bars represent mean ± SEM (n = 4). *p ≤ 0.05, **p ≤ 0.01, ***p ≤ 0.001, ****p ≤ 0.0001, Two-way Anova, Tukey’s multiple comparison test, (n=4). Cell illustrations created in BioRender by E.M. Aksu. https://BioRender.com/xhsb61e

### AAG and TDG exert antagonistic control over synaptic gene expression

Loss of AAG primarily leads to preterm activation of gene expression during neuronal differentiation (Fig. 2). Similarly, the DNA glycosylase TDG has been shown to be essential for the regulation of gene expression and neuronal fate^9^, suggesting that DNA glycosylases may have a broader role in controlling neuronal genes. To investigated if AAG and TDG share target genes, mRNA sequencing (RNA-seq) was performed on WT and sgAAG hiPSCs (D0) and D4, D7 and D14 iNs. The resulting datasets were integrated with the previously published sgTDG iNs data for comparative analysis^9^. This led to identification of 380 genes commonly deregulated across all differentiation timepoints (D4 to D30) in both sgAAG and sgTDG iNs, compared to WT samples and D0 (Fig. 3a). Notably, clustering of the differentially expressed genes (DEGs) indicated that AAG and TDG regulate expression in the opposing manner. Subsequent gene ontology analysis of Cluster 1, encompassing 51 most significant DEGs whose expression is upregulated in sgAAG and downregulated in sgTDG iNs, revealed synaptic processes including: sodium, calcium and potassium transport, as well as synaptic transmission, being the most significantly affected (Fig. 3d). These findings were further supported by expression analysis of key DEGs, *SLC17A7* encoding vesicular glutamate transporter 1 (vGLUT1), and synaptic *PDLIM4* and *SLC6A3* genes, all of which were significantly upregulated in sgAAG and downregulated in sgTDG iNs (Fig. 3b, c and Supplementary Fig. 3a-d). In summary, these results suggest that AAG and TDG exhibit antagonistic control over gene expression in iNs. To further test if this effect could be mediated through direct crosstalk between the two DNA glycosylases, proximity ligation analysis (PLA) was performed. Interestingly, AAG and TDG localized in very close spatial proximity, less than 40 nm, in the nuclei of iNs and hiPSCs (Fig. 3e and Supplementary Fig. 3e). The obtained PLA foci were specific, as no signal was detected in the control iNs lacking AAG (sgAAG). Together, these results suggest that AAG and TDG could engage in functional crosstalk within nuclei of developing neurons, of relevance for gene expression regulation.

**Figure 3.**
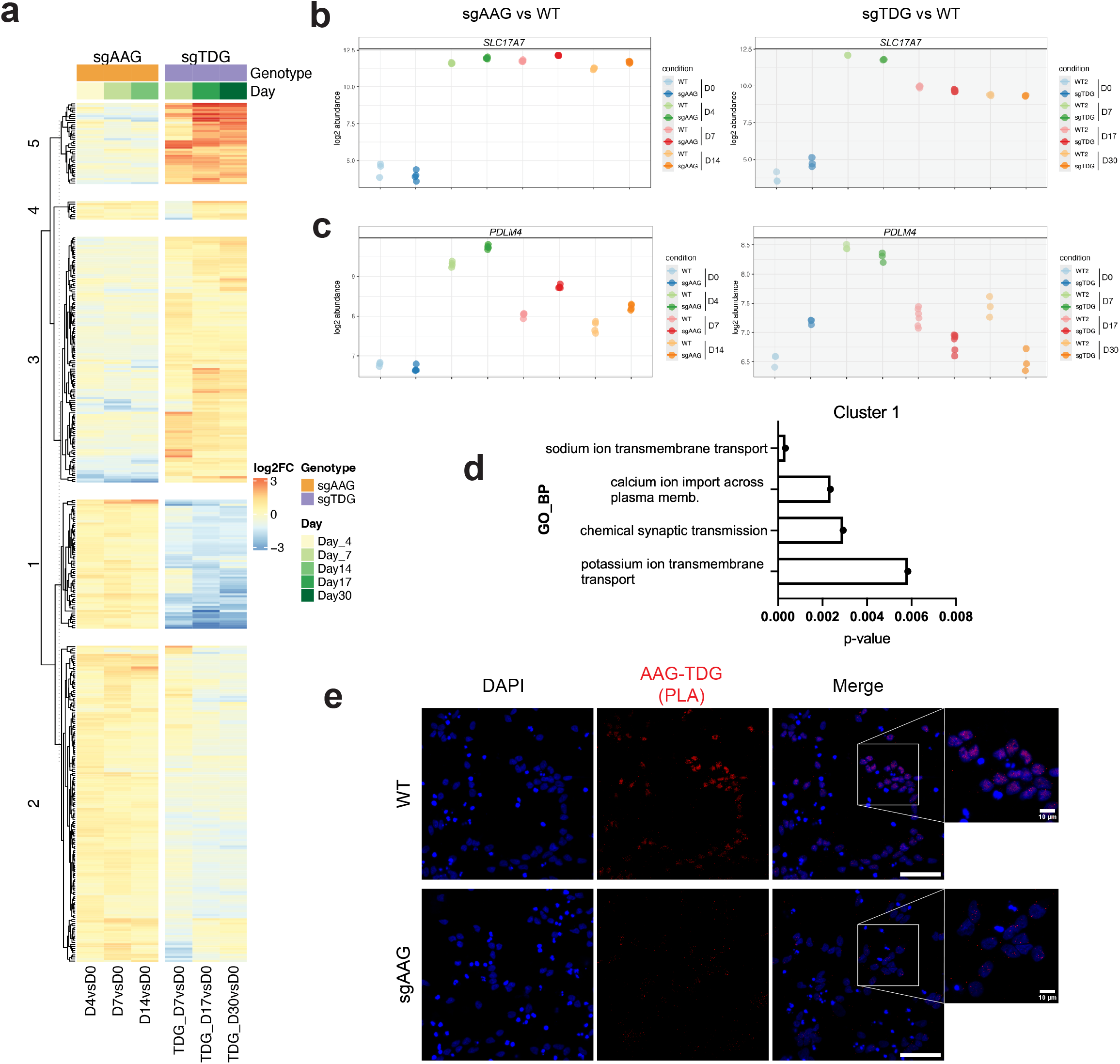
Synaptic gene expression is opposingly cross-regulated by AAG and TDG. **a** Heatmap of differentially expressed genes (DEGs) in AAG knock-down (sgAAG) vs wild type (WT), and sgTDG vs WT over the course of neuronal differentiation; day (D) 4 to D30. DEGs were clustered into 5 distinct groups using consensus k-means, color scale represents fold change. **b-c** Relative gene abundance of cluster 1 marker genes *SLC17A7* (c) and *PDLIM4* (d) during neuronal differentiation in sgAAG vs WT in the left, and sgTDG vs WT in the right panels **d** Top four most significant biological processes (BP) gene ontology (GO) terms, identified by DAVID, for genes from cluster 1, presented in (a). **e** Proximity Ligation Assay (PLA) showing AAG and TDG in the close proximity, in D7 iNs (Scale bar 50 µm, scale bars magnified images = 10 µm).

### By inhibiting TDG glycosylase activity AAG regulates global levels of oxidized methylcytosines and neuronal DNA breaks

TDG regulates developmental gene expression in differentiating cells by initiating BER and subsequently generating SSB intermediates^9, 37^. Moreover, we showed that AAG status impacts endogenous double strand DNA breaks (DSBs) marked by γH2AX in iNs (Fig. 1h, i) and mouse hippocampus^29^. In line with this and considering the opposing impacts of AAG and TDG on neuronal gene expression (Fig. 3), we next investigated global DNA damage levels (SSBs and DSBs) in AAG lacking (sgAAG) iNs by comet assay (Fig. 4a). Notably, sgAAG iNs showed higher levels of DNA damage compared to WT at early stages of differentiation (D4 and D7), whereas this difference became non-significant in more mature D14 neurons (Fig. 4b). To further confirm that the increase in DNA damage levels is due to accumulation of SSBs in addition to DSBs, poly(ADP-ribose) (PAR) levels were analyzed by immunoblot analysis in D7 iNs (Fig. 4c, d) and D0 stem cells (Supplementary Fig. 4a, b). PAR levels were significantly increased in D7 sgAAG iNs, but not stem cells, suggesting that SSBs specifically accumulate in developing neurons lacking AAG. Since TDG-driven removal of oxidized methylcytosines (e.g., 5fC) is a main driver of DNA breaks in differentiating iNs, the elevated SSB levels in sgAAG cells imply that AAG might act as a negative regulator of TDG activity. To test this, we performed biochemical analysis of TDG glycosylase activity on 5fC containing DNA in the presence of increasing AAG concentrations, or BSA as a control (Fig. 4e). Importantly, presence of AAG resulted in significant reduction of TDG-mediated 5fC excision, while BSA at equivalent concentrations had no effect (Fig. 4f). These results suggest that AAG inhibits TDG activity on 5fC. To investigate whether this effect takes place in cells, liquid chromatography-tandem mass spectrometry (LC-MS/MS) analysis was performed next. Interestingly, while no differences in 5mC and 5hmC status were observed between the genotypes (Supplementary Fig. 4c, d), levels of 5fC and 5caC were significantly reduced in sgAAG compared to WT cells (Fig. 4g, h). This further suggests that in the absence of AAG, TDG activity is increased, resulting in higher removal of oxidized methylcytosines, consistent with increased breaks and PAR levels detected in sgAAG cells. The observed changes in 5fC and 5caC status were not due to differences in TDG levels, as TDG mRNA expression remained unchanged in sgAAG, compared to WT cells (Supplementary Fig. 4e). Taken together, our results identify AAG as a regulator of active DNA demethylation and neuronal DNA break formation through modulation of TDG activity.

**Figure 4.**
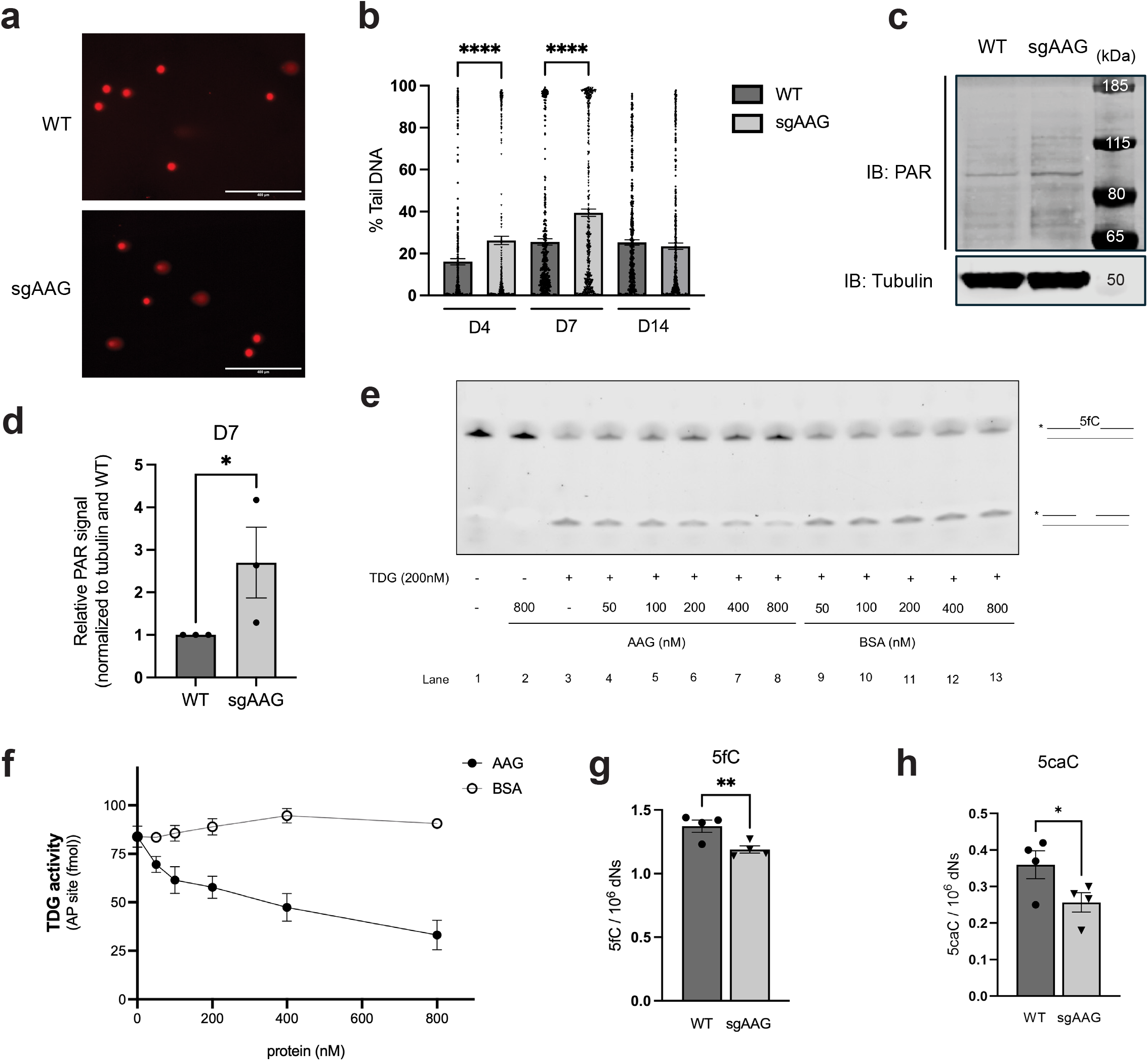
Lack of AAG promotes TDG’s catalytic activity, leading to decrease in global levels of oxidized methylcytosines and increase in DNA breaks of differentiating neurons. **a** Representative comet assay images of wild type (WT) and AAG knock-down (sgAAG) iNs at D7. Scale bar 400 µm. **b** Analysis of comet assays (as the one in a) showing global DNA damage levels in WT and sgAAG iN at D4, D7 and D14. Error bars represent mean ± SEM (n=6 coverslips, from 2 independent experiments), one-way Anova, Tukey’s multiple comparison test, ****p ≤ 0.0001. **c** Representative images of immunoblot analysis targeting poly ADP ribose (PAR) in WT and sgAAG at D7 iNs. Tubulin serves as loading control. **d** Quantification of PAR levels (from n=3 experiments as the one in (c)) Error bars represent mean ± SEM, one-tailed, Mann-Whitney test. *p ≤ 0.05. **e** Representative DNA Glycosylase Assay showing cleavage activity of 200 nM TDG on 5fC containing DNA oligonucleotide in the presence of AAG at varying concentrations; with BSA used as a control. **f** Quantification of three independent experiments as the one in (e). The bars depict mean ± SEM. **g-h** LC-MS/MS analysis of oxidized methylcytosines 5-formylcytosine (5fC) (g) and 5- carboxycytosine (5caC) (h); n=4, one-tailed Student’s t-test *p ≤ 0.05, **p ≤ 0.01.

### Single-molecule resolution of AAG binding to 5fC, which results in reduced TDG affinity for the shared substrate

AAG could impair TDG activity either through direct interaction with TDG or through substrate competition by binding 5fC and thereby limiting TDG access. To identify the mechanism through which AAG alters TDG activity, single molecule analysis of DNA-binding proteins from nuclear extracts (SMADNE)^38^ was conducted using C-Trap (Lumicks). This approach determines the single-molecule kinetics of DNA glycosylase binding. In brief, the method involves capturing streptavidin-coated beads with optical tweezers and subsequent tethering of biotinylated DNA. Upon equilibration, imaging is initiated as DNA is brought into HEK293T nuclear extracts of either transiently overexpressed TDG-HaloTag, AAG-EGFP, or a mixture of the two. (Fig. 5a). Nuclear extracts were generated from HEK293T cells transiently overexpressing JF-635 labelled TDG-HaloTag or AAG-EGFP. Characterization of the overexpressed TDG-HaloTag and AAG-EGFP with SDS-PAGE revealed that the full-length proteins were the primary species within the extracts (Supplementary Fig. 5a, b). The DNA template was designed to contain a single 5fC modification in the middle of a 12.6kb construct, through ligation of 30 nt 5fC-containing double-stranded DNA oligonucleotide into 6.3kb DNA handles (Lumicks) (Fig. 5b). The kymographs of specific DNA glycosylase binding to the 5fC- containing DNA appeared as straight horizontal lines in the middle of the template (Fig. 5b), where Y coordinate defines DNA binding position and X coordinate depicts the duration of binding events (dwell time). Additionally, time between two binding events is defined as gap time. Sorting of dwell times and gap times by length yielded a cumulative residence time distribution (CRTD) and cumulative gap time distribution (CGTD), respectively. Fitting of CRTD for TDG to a double-exponential decay function identified 80% of short 5fC binding events (0.9 ± 0.03 s) and 20% long 5fC binding events (12.6 ± 2.8 s), compared to an average 5fC- nonspecific binding time of 0.84 ± 0.005 s (Fig. 5c, Table 1). The measured 5fC binding times are comparable with the previous findings^39^. Similarly, gaps between events were classified by length and represented as CGTD and knowing the concentration of TDG in the flowcell resulted in a TDG on rate of 7.1 ± 0.7 x 10^8^ s^-1^ M^-1^ (Fig. 5c, Table 1). The single-molecule derived equilibrium dissociation constant (*K_D_*) was determined next as the ratio of the off- and the on-rate, resulting in 0.7 ± 0.1 nM, thus indicating high TDG affinity for 5fC sites.

**Figure 5.**
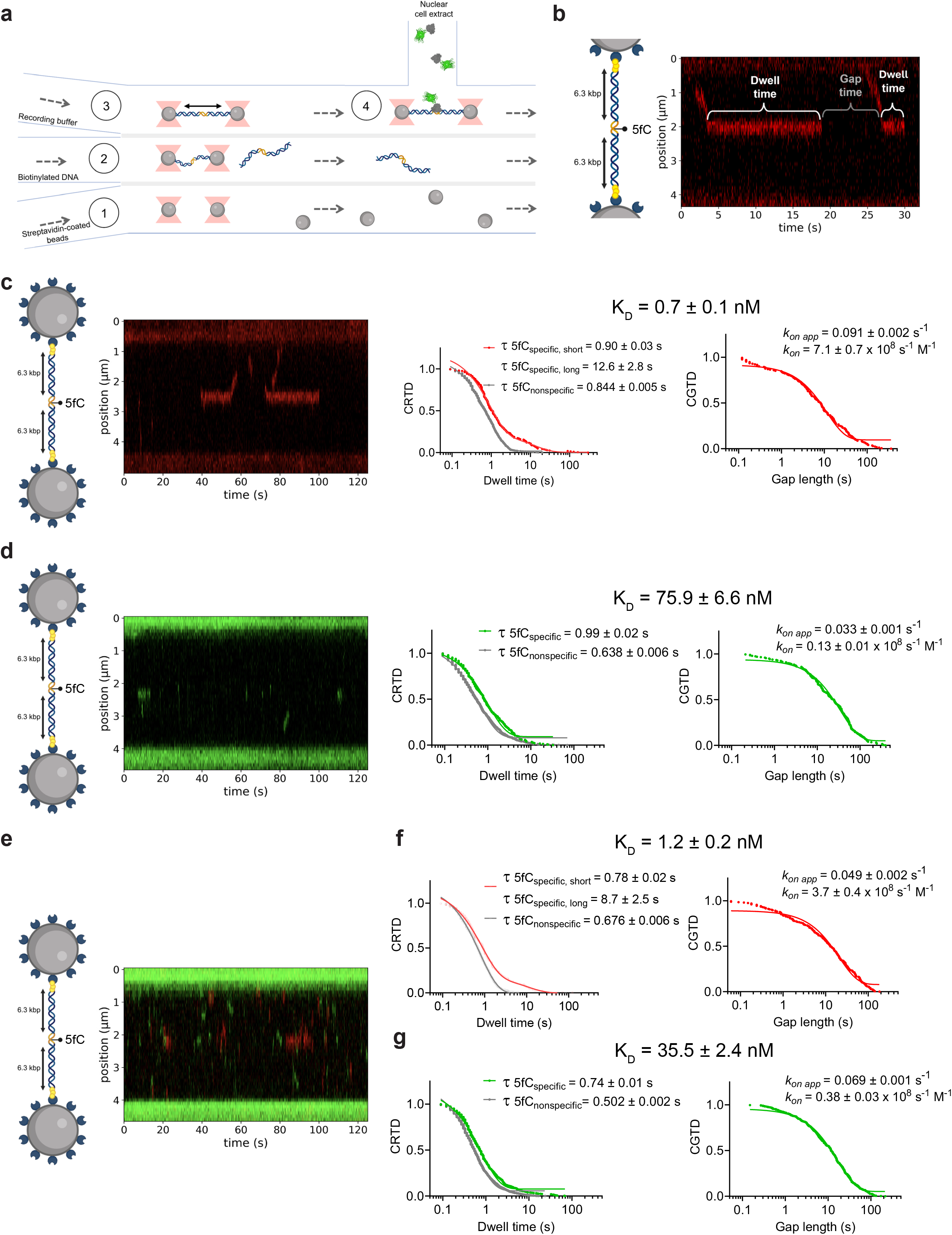
At the single-molecule level AAG binds to 5fC simultaneously reducing TDG affinity for the shared substrate. **a** Schematic representation of the experimental workflow. Upon optical trapping of the streptavidin-coated beads in channel 1; 12.6kb DNA with a single 5-formylcytosine (5fC) positioned in the middle of the template is tethered between the beads in channel 2; and substrate washing in channel 3; binding of the fluorescently labelled overexpressed TDG-HaloTag and/or AAG-EGFP is imaged in channel 4 containing nuclear extract. **b** Representative kymograph data depicting TDG-HaloTag (red) engaging with 5fC in the middle of 12.6-kbp tethered DNA (grey lines are stationary events), and an example of dwell/gap times (white lines) utilized to determine on and off rates. **c-g** Representative kymograph data of TDG-HaloTag (in red) (c, e) and AAG-EGFP (in green) (d, e) binding to 5fC; with cumulative residence time distribution (CRTD) and cumulative gap time distribution (CGTD) plots shown on the right. The CRTD analysis used to estimate dwell time distribution following numbers of binding events obtained from three independent experiments were included: (c) TDG alone, n=207 5fC specific events and n=742 5fC non-specific events; (d) AAG alone, n=230 5fC specific events and n=834 5fC non-specific events; (f) TDG in the presence of AAG, n=195 5fC specific events and n=627 5fC non-specific events; (g) AAG in the presence of TDG, n=608 5fC specific events and n=2093 5fC non-specific events. The observed CRTD, used to determine the apparent 5fC association rate, and the CGTD, used to identify the on rate for binding, were combined to determine *K_D_* values for interactions (reported in Table 1). Experimental c-trap workflow and DNA figures created in BioRender by Gjøsæther, A. https://BioRender.com/6r4aa8y

**Table 1.**
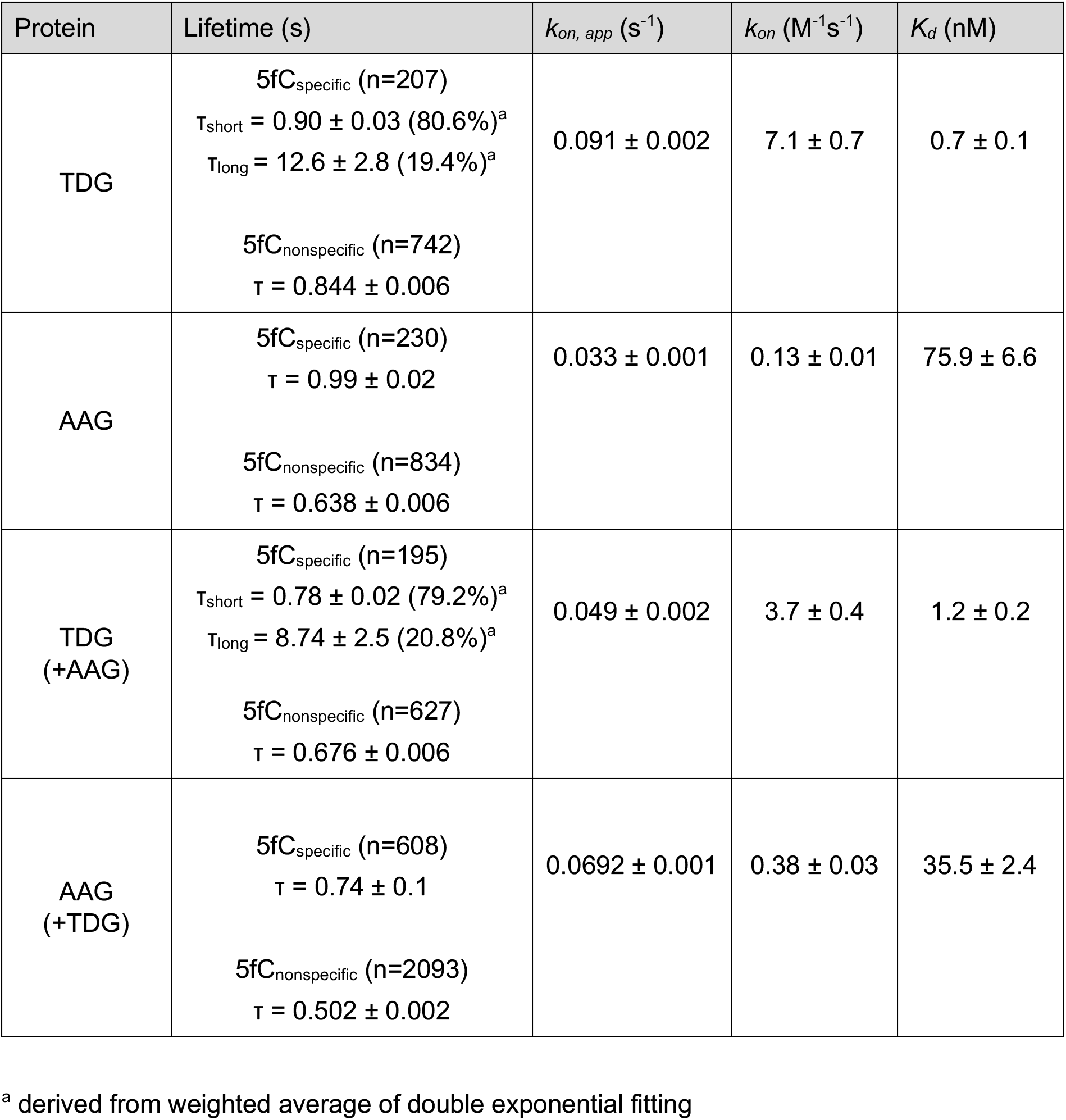
Binding kinetics together with the errors across the datasets.

| Protein | Lifetime (s) | *k*<sub>*on, app*</sub> (s<sup>-1</sup>) | *k*<sub>*on*</sub> (M<sup>-1</sup>s<sup>-1</sup>) | *K*<sub>*d*</sub> (nM) |
|---|---|---|---|---|
| TDG | 5fC<sub>specific</sub> (n=207)<br>τ<sub>short</sub> = 0.90 ± 0.03 (80.6%)<sup>a</sup><br>τ<sub>long</sub> = 12.6 ± 2.8 (19.4%)<sup>a</sup><br>5fC<sub>nonspecific</sub> (n=742)<br>τ = 0.844 ± 0.006 | 0.091 ± 0.002 | 7.1 ± 0.7 | 0.7 ± 0.1 |
| AAG | 5fC<sub>specific</sub> (n=230)<br>τ = 0.99 ± 0.02<br>5fC<sub>nonspecific</sub> (n=834)<br>τ = 0.638 ± 0.006 | 0.033 ± 0.001 | 0.13 ± 0.01 | 75.9 ± 6.6 |
| TDG<br>(+AAG) | 5fC<sub>specific</sub> (n=195)<br>τ<sub>short</sub> = 0.78 ± 0.02 (79.2%)<sup>a</sup><br>τ<sub>long</sub> = 8.74 ± 2.5 (20.8%)<sup>a</sup><br>5fC<sub>nonspecific</sub> (n=627)<br>τ = 0.676 ± 0.006 | 0.049 ± 0.002 | 3.7 ± 0.4 | 1.2 ± 0.2 |
| AAG<br>(+TDG) | 5fC<sub>specific</sub> (n=608)<br>τ = 0.74 ± 0.1<br>5fC<sub>nonspecific</sub> (n=2093)<br>τ = 0.502 ± 0.002 | 0.0692 ± 0.001 | 0.38 ± 0.03 | 35.5 ± 2.4 |

<sup>a</sup> derived from weighted average of double exponential fitting

Subsequent analysis of AAG binding identified an average binding time of 0.99 ± 0.02 s to 5fC compared to 0.638 ± 0.006 s on unmodified DNA, indicating a preference of AAG for the 5fC compared to unmodified DNA (Fig. 5d, Table 1). To our knowledge, these results represent the first direct visualization of AAG engagement with 5fC at the single-molecule level (Fig. 5d) and further support previous multi-omics and biochemical studies that identified AAG as reader of 5fC^24, 25^. Analysis of the CGTD identified an on rate of 0.13 ± 0.01 x 10^8^ s^-1^ M^-1^ and in combination with binding times measurements yielded a *K_D_* of 75.9 ± 6.6 nM for AAG on 5fC (Fig. 5d). The identified AAG dissociation rates are of the same order of magnitude as very recently reported dissociation constants of AAG binding to 5fCs in the promoter sequence of mouse *mSP1* determined by electromobility shift analysis^40^. Taken together, these results indicate that AAG efficiently binds 5fC, though with a lower affinity than TDG.

Next, to investigate if AAG presence impacts the binding kinetics of TDG to 5fC, the nuclear extracts were mixed before adding to the flow cell (Fig. 5e). While the AAG presence did not majorly affect short 5fC binding times, it reduced long TDG binding to 5fC to 8.7 ± 2.5 s (Fig. 5f, Table 1). Notably, presence of AAG had the most prominent effect on the TDG on rate, resulting in 3.7 ± 0.4 x 10^8^ s^-1^ M^-1^, (Fig. 5f, Table 1), nearly two-fold decrease in comparison to the condition without AAG (Fig. 5d, Table 1). The change was further reflected in a decrease in TDG affinity for 5fC, resulting in *K_D_* = 1.2 ± 0.2 nM (Fig. 5f, Table 1). This suggests that AAG reduces TDG affinity for 5fC. Interestingly, presence of TDG exhibited opposite effect on AAG (Table 1). While AAG dwell times were largely unchanged in the presence of TDG, the on rate of AAG was increased by nearly three-fold and resulted in reduced *K_D_* of 35.5 ± 2.4 nM (Fig. 5g, Table 1). We hypothesize that TDG transiently alters DNA topology upon binding (e.g. by DNA bending and minor groove widening^41^), thereby creating a local environment that facilitates faster AAG binding. This is further supported by an overall increase in the number of AAG binding events in the presence of TDG (Table 1).

To further determine whether these kinetic changes arise from colocalization or competitive binding of the proteins to the shared substrate, colocalization assessment was conducted (Supplementary Fig. 5c, d). Only limited colocalization between AAG and TDG was detected, with the predominant event being binding of a single DNA glycosylase (AAG or TDG) to 5fC (Supplementary Fig. 5c). Subsequent analysis of colocalization events as defined in ^38^, revealed that when AAG and TDG do bind 5fC together, the colocalization occurs in a way that one DNA glycosylase binds first, followed by second DNA glycosylase binding and dissociation, after which the initially bound DNA glycosylase dissociates (category 5 and 9 events) (Supplementary Fig. 5d). Together, these results indicate that the presence of AAG alters TDG binding kinetics to 5fC sites primarily through competitive substrate binding.

## Discussion

In this study, we investigated the mechanism through which two DNA glycosylases temporally regulate gene expression during neuronal development, by using approaches across scales from brain tissue analysis to single molecule visualization of DNA glycosylases competition for shared substrates. The primary role of DNA glycosylases is widely considered to be maintenance of genome integrity under exogenous and endogenous insults^42^. In addition to these canonical functions, mounting evidence indicates that DNA glycosylases play important developmental roles of particular importance for cognition and behavior^9, 29–33^. In line with this, we recently showed that AAG loss results in reduced anxiety and improved memory in adult mice^29^. Despite the emerging unconventional roles, the detailed mechanistic insights into how DNA glycosylases directly affect brain functions are largely missing. In this work, we demonstrate that AAG deficiency adversely affects neurogenesis (Fig. S1). Similarly, loss of another DNA glycosylase, NEIL3, was reported to adversely affect NSC/NPC properties^43, 44^, thus suggesting a broader DNA glycosylase phenomenon. Interestingly, single-cell RNA sequencing analysis of iNs across differentiation stages indicates that AAG deficiency alters temporal regulation of gene expression (Fig. 2a), likely contributing to the observed impairment in neurogenesis. Preterm activation of synaptic genes and impaired neurodevelopmental gene expression (Fig. 2d, e) suggest premature differentiation in AAG deficient cells. Reduced expression of OGG1 and NEIL3, DNA glycosylases involved in repair of oxidative DNA damage, was similarly indicated to impair neurodevelopmental gene expression^44^. Together with the previously established role of TDG in differentiation^9^, these findings imply a potential global importance of DNA glycosylases in the temporal regulation of developmental gene expression.

Regulation of developmental gene expression has been shown to be dependent on active demethylation driven by BER. During this, SSB intermediates are created in enhancer and promoter regions of neuronal genes^10, 11^, through TDG action on oxidized methylcytosines in active DNA demethylation^9^. Besides TDG, AAG is the only other DNA glycosylase suggested to directly bind these modifications^24, 25, 40^. By using a single molecule approach, we provide direct visualization of AAG binding to 5fC (Fig. 5). While AAG as expected has lower affinity than TDG for 5fC (Table 1), the identified single molecule dissociation constant (*K_D_* of 75.9 ± 6.6 nM, Fig. 5d) is comparable to recently reported AAG binding to 5fC in context of mouse *SP1* promoter^40^. Notably, AAG appears to potentially bind 5fC with comparable affinity to that of its cognate substrate hypoxanthine^45^. Together, these findings further support the notion that AAG functions as reader of 5fC. Though single molecule analysis has been performed to assess the binding kinetics of AAG and TDG DNA glycosylases to their cognate substrates previously^39, 46^, this study is the first to investigate how the binding kinetics of these two proteins are altered when engaging the shared modification within the same nuclear extract. In the presence of AAG, the affinity of TDG is reduced (Fig. 5f, Table 1), likely due to competitive binding of two glycosylases for 5fC rather than through their colocalization at 5fC (Supplementary Fig. 5). In line with this, and together with the observed increase in the levels of DNA breaks upon AAG loss (Fig. 1h, i and 4a, b), we propose a model in which DNA glycosylases regulate active DNA demethylation during neurodevelopment (Fig. 6). In the WT cells, AAG and TDG compete for the shared 5fC substrates, thereby fine-tuning the DNA demethylation process, associated DNA break induction, and neuronal gene expression. In the absence of AAG, TDG has increased affinity for 5fC sites, resulting in accumulation of single-strand DNA breaks and preterm expression of TDG regulated synaptic genes (Fig. 3 and ^9^), ultimately impairing neuronal development. In addition, accumulating evidence indicates complex interactions between synaptic functions and genome dynamics, especially DNA repair associated proteins, both in basal state and neuronal activity dependent manner^47–49^, with several participating in active demethylation^50^. However, future work is needed to investigate the impact of DNA glycosylase crosstalk in context of neuronal activity and functional synaptic transmission.

**Figure 6.**
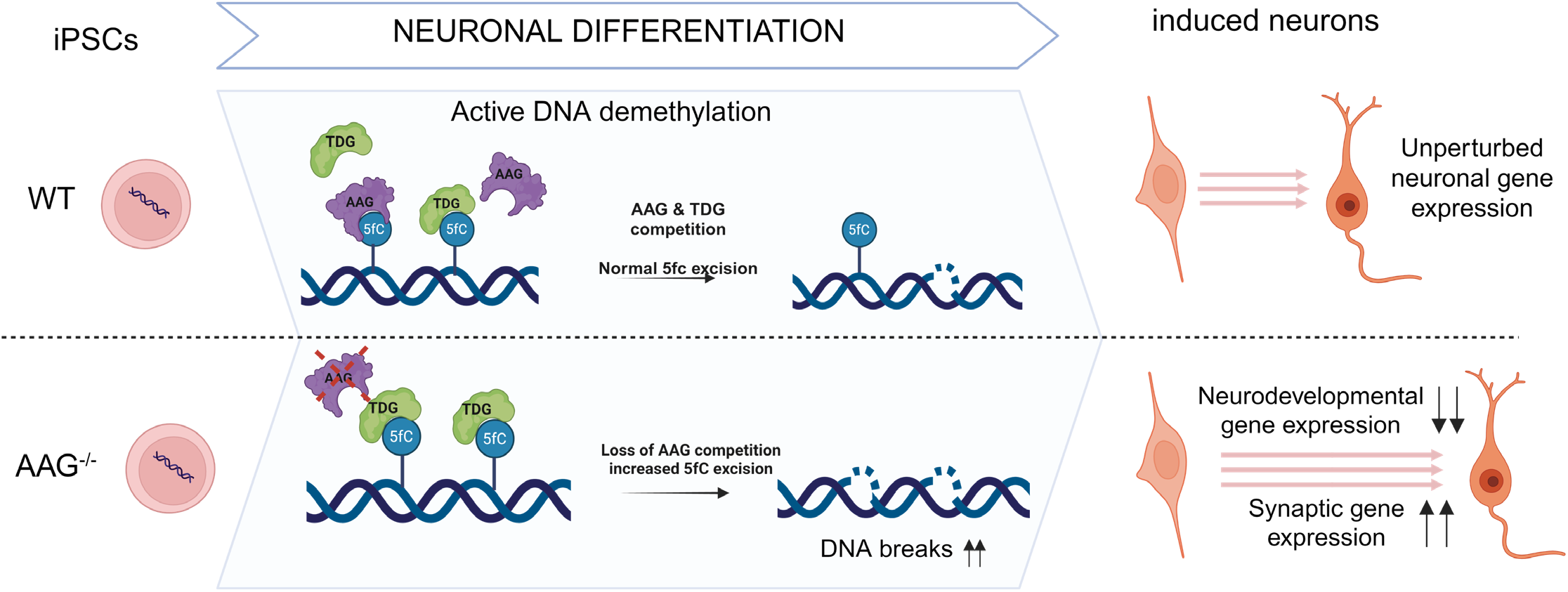
Model for the DNA glycosylases regulated neurodevelopmental gene expression through substrate competition during active DNA demethylation. In wild type (WT) cells alkyladenine DNA glycosylase (AAG) and thymine DNA glycosylase (TDG) compete for binding to shared 5-formylcytosine (5fC) sites. The competitive binding fine-tunes TDG-mediated 5fC excision during DNA demethylation process, the associated DNA break induction, and consequently neuronal gene expression. In the absence of AAG, TDG has increased affinity for 5fC sites, which results in: increased excision of 5fC during active demethylation, elevated levels of DNA breaks, altered temporal expression of neurodevelopmental and synaptic genes, and consequently impaired neurodevelopment. Schematic representation was created in BioRender by Aksu. E.M https://BioRender.com/lutj2cj

We report that lack of AAG in differentiating iNs, besides the increase in SSB, also contributes to increase in DSB (Fig. 1h, i and 4a, b). Since CpGs are symmetrically methylated in mammals, it is possible that during demethylation TDG concomitantly removes modifications at the opposite DNA strands, thereby giving rise to clustered SSBs and consequently increased DSB levels in sgAAG cells.

Despite TDG and AAG being the direct binders of oxidized methylcytosines, several other DNA glycosylases are shown to impact active demethylation. NEIL1 and NEIL2 were demonstrated to promote TDG-mediated removal of 5fC and 5caC^51^. NEIL1 was additionally shown to excise 5caC and enhances TDG activity on 5fC and 5caC^52^. UNG2 was similarly shown to impact 5fC and 5caC levels in association with TET2, although the mechanism of this reduction remains unknown^53^. While current understanding of the coordinated activity of DNA glycosylases at the epigenetic modifications remains limited, the findings of this study, together with accumulating evidence supporting critical roles of DNA glycosylases in development, highlight the need for further investigations into glycosylase functions in active DNA demethylation.

Taken together, we identify the coordinated mechanism of AAG and TDG action at the 5fC sites that determines DNA breaks dynamics and consequently temporal regulation of gene expression, needed for unperturbed neuronal development. As such this work provides novel evidence for the functional importance of DNA glycosylase competition at shared substrates, extending beyond single–DNA glycosylase–driven pathways and the traditionally hypothesized enzyme backup roles.

## Methods

### Mice and genotyping

C57BL/6J wild type (WT) and Aag knock-out *Aag*^-/-^ mice, kind gifts from prof. Leona D. Samson (MIT, USA)^54^, were maintained in pathogen-free conditions at the Comparative Medicine Core Facility of the Norwegian University of Science and Technology (NTNU). The colonies underwent backcrossing for 3 generations before the start of experiments. To reduce genetic drift, both *Aag*^-/-^ mice were regularly backcrossed after 7–10 generations. The experiments were approved and conducted in accordance with the Norwegian Animal Research Authority. WT and *Aag*^-/-^ were genotyped according to^54^. Sex determination of pups at postnatal day five (P5) was conducted by the amplification of the Y chromosome gene *Sry* by PCR as described previously^29^.

### Isolation and propagation of Neural Progenitor Cells (NPCs)

Brains from newborn (P0-P1) WT and *Aag^-/-^*mice were isolated, cut in pieces and mechanically dissociated by pipetting, to obtain single cells. The cell suspension was transferred to a 25 cm^2^ culture flask containing 7 ml proliferation media (DMEM/F12 with glutamine (Gibco, 11320033), 1% penicillin/streptomycin (Sigma, P4333), 1% N2 supplement (ThermoFisher Scientific, 17502048), 2% B-27 supplement minus vitamin A (Gibco, 12587010), 10 ng/ml of basic human FGF (R&D Systems, 233-FB-010) and 20 ng/ml of human EGF (R&D Systems, 236-EG-200)). After 7 days, the resulting neurospheres were trypsinized, washed and propagated. All experiments were conducted between passages 3 to 8. Two independent clones from WT and *Aag^-/-^* NPCs were used for the experiments.

### Proliferation, self-renewal and differentiation of NPCs

To evaluate the proliferation capacity, neurospheres were trypsinized, counted and plated at a density of 8,000 cells/well of a 96 well plate. Cell proliferation was evaluated after 1 h (day 0) and next every 24 h during five days by using PrestoBlue Cell Viability Reagent (Invitrogen, A13262). Absorbance was measured by FLUOstar Omega (BMG Labtech). The experiment was repeated three times including 6 technical replicates per clone. For the assessment of the self-renewal, 10,000 single cells were plated per well of a 6-well plate and grown for 10 days. Images from spheres forming cells were captured in bright field using the tile scan mode of EVOS FL Auto microscope (ThermoFisher Scientific) with 4x objective. All neurospheres (>50 µm) were counted using Fiji software v. 1.54f (National Institutes of Health). Results represent the number of spheres forming cells relative to WT from four independent experiments, each including two technical duplicates.

For differentiation of NPCs, cells were plated in 48 wells plates pre-coated with poly-D-lysine (Gibco, A38 90401) at a density of 50,000 cells/well. Differentiation was induced with proliferation media containing 2% B-27 supplement with vitamin A (Gibco, 17504001), with daily media exchanges. After five days, differentiated NPCs were subjected to immunofluorescence analysis.

### CRISPRi Plasmids

Non-targeting (Ng) and AAG targeting sgRNA (Supplementary Table 1) oligonucleotides were annealed at 95 °C, 5 min and gradually cooled to room temperature (RT). Plasmid pU6-sgRNA EF1Alpha-puro-T2A-BFP (Addgene: #60955)^55^ was digested with BstXI (R0113S, NEB) and BlpI (R0585S, NEB). Annealed sgRNAs were next ligated into pU6-sgRNA EF1Alpha-puro- T2A-BFP with T4 ligase (M0202S, NEB) at 16°C overnight. Competent DH5α *E. coli* were incubated 30 min with respective plasmids on ice, and heat shock transformed at 42 °C for 45 sec, 1min on ice. Bacteria were recovered for 1 h at 37 °C, seeded on LB agars containing ampicillin and incubated overnight at 37 °C. Single colonies were next propagated in LB medium containing ampicillin and plasmids were isolated with Miniprep (Wizard® *Plus* SV, Promega) according to manufacturer’s instructions. Plasmids were Sanger sequenced to confirm the constructs.

### Generation of Lentivirus for CRISPRi

Human embryonic kidney (HEK293T) cells were grown in DMEM high glucose media (Sigma, D6429) with 10% fetal bovine serum (Sigma, F7524), 1 % penicillin/streptmycin (Gibco, 15140122), and maintained in humidified 5% CO_2_ atmosphere, at 37 °C. For each virus, packaging plasmids psPAX2 (Addgene: #12260) and pMD2.G (Addgene: #12259), and pU6- sg-Ng EF1Alpha-puro-T2A-BFP or pU6-sg-AAG EF1Alpha-puro-T2A-BFP were mixed in 2:1:2 with CaCl_2_ and HEPES buffered saline. psPAX2 and pMD2.G were gifts from Didier Trono. Mixtures were incubated at RT for 5 min and added to HEK293T cells. Next day, cells were washed, medium exchanged, and virus were collected 72 hr after transfection.

### Stem cell culture

The human induced pluripotent cells (iPSCs) WTC11 used in this study previously engineered to express CAG-dCas9-BFP-KRAB ^56^ and NGN2 under inducible promoter ^34, 35^. The cells were cultured on Geltrex (Gibco, A1413202) coated plates in Essential 8 (E8) medium (Gibco, A1517001). The cells were cultured in a humidified 5% CO_2_ atmosphere at 37°C. The cells were regularly tested for mycoplasma and karyotyped (StemCell Technologies, # 07550).

### Viral transduction

WTC11 iPSCs were dissociated with accutase (StemCell Technologies, 07920), neutralized with DMEM/F12 (Gibco, 11330032), pelleted at 200 x g and counted. 1 x 10^6^ stem cells were transduced in well of a 6-well plate with non-targeting or AAG sgRNA expressing lentivirus in E8 medium containing ROCK Inhibitor (ROCKi) (Calbiochem, 688000) and polybrene. Medium was removed next day and cells washed with PBS. After 24 h cells were split with accutase and seeded at low density in E8 medium with ROCKi and 10 µg/ml puromycin. Cells were next day washed with PBS and grown in E8 with Rocki for 24 h. Medium was then changed into regular E8 for another 24 h.

### Induced Neuron (iN) generation

On day one (D1) 5 x 10^6^ iPSCs were seeded on 10 cm Geltrex coated dish in N2 medium ((DMEM)/F12 (Gibco, 11330032), N2 supplement (Gibco, 17502048), GlutaMAX (Thermofisher Scientific, 35050061), MEM non-essential amino acids (NEAA) (Thermofisher Scientific,11140050), 10 μM ROCKi (Y-27632;), and 2 μg/ml doxycycline (Merck, D9891-5G)) with daily media changes. On D3, cells were dissociated with accutase and reseeded to PLO (Sigma, P4957) and Laminin (R&D Systems 3446-005-01) coated plates in BrainPhys medium (StemCell Technologies, 05790) supplemented with B27 Plus supplement (Gibco, 17504001), NT-3 (Peprotech AF-450-03), BDNF (R & D Systems, 248-BD) and Laminin (R&D Systems, 3446-005-01). Half medium was changed on D6, followed by medium changes every second day. iNs were collected at D4, D7 and D14 for the subsequent analysis.

### Gene expression analysis

Total RNA was isolated with RNeasy mini kit (Qiagen,74106) according to manufacturer’s protocol and reverse transcribed with high-capacity cDNA RT kit (ThermoFisher, 4368814). The quantitative PCR analysis was performed on StepOnePlus Real-Time PCR system, with Power SYBR Master Mix 2X (Applied Biosystems, 4367659), cDNA and target forward and reverse primers, listed in Supplementary Table 2. Relative transcript levels were determined by normalizing to ACTB mRNA levels.

### Alkaline Comet FLARE

iNs were collected with accutase, counted, mixed with 1% low melting point agarose (Lonza, 50100) and applied onto glass slides (Trevigen). Cells were lysed for 1 h at 4 °C with lysis buffer (1% Triton, 10% DMSO, 1% sodium lauryl sarcosinate, 100 mM EDTA, 10 mM Tris, 2.5 M NaCl) and washed 3 x 10 min in FLARE buffer (25 mM HEPES-KOH at pH7.4, 0.25 M KCl, 25 mM EDTA pH8). Next, the slides were incubated with either AAG (NEB, M0313S) or AAG storage buffer (negative control) in FLARE reaction buffer for 30 min at 37 °C. After incubation, the slides were washed twice for 10 min with alkali unwinding solution (200 mM NaOH, 1 mM EDTA, pH > 13) and subjected to 20 min electrophoresis in alkali unwinding solution at 25V, 300 mA. Slides were washed with water 2 x 10 min and finally with 70% ethanol for 5 min. Upon drying 30 min at 37 °C, 1:10,000 diluted SYBR Green (Invitrogen, 57563) solution was added and the slides imaged using EVOS FL Auto with 10x objective. Quantifications were performed using Comet Assay IV Software (Instem, Perceptive Instruments)

### LC-MS/MS analysis of DNA modification

DNA was extracted using the DNeasy kit (Qiagen, 69506) and digested to deoxynucleosides with nuclease P1 (Sigma, N8630), benzonase (Santa Cruz Biotech, sc-202391) and alkaline phosphatase (Sigma, P5931) in 10 mM ammonium acetate pH 6.0 and 1 mM magnesium chloride at 40 °C for 1 h. Digested samples were precipitated with 3 volumes of acetonitrile and supernatants were lyophilized and dissolved in a solution of internal standards (I.S.) for analysis. An Agilent 1290 Infinity II UHPLC system with an ZORBAX RRHD Eclipse Plus C18 150 x 2.1 mm (1.8 μm) column protected with an ZORBAX RRHD Eclipse Plus C18 5 x 2.1 mm (1.8 µm) guard column (Agilent) was used for chromatographic separation. The mobile phase consisted of A: water and B: methanol (both added 0.1% formic acid), for 5hm(dC), 5f(dC) and 5ca(dC) starting at 0.15 ml/min flow of 5% B for 1 min followed by 3.5 min gradient of 5-15 % B, 3 min of 15-95% B while increasing the flow to 0.23 ml/min, 0.5 min of 95% B, and 4 min re-equilibration with 5% B. Unmodified nucleosides and 5m(dC) were chromatographed at 0.23 ml/min with a 4 min gradient 5-95% B and 4 min re-equilibration with 5% B. Mass spectrometric detection was performed using an Agilent 6495 Triple Quadrupole system operating in positive electrospray ionization mode. The following mass transitions were monitored: 242.1/126.1 (5m(dC)); 258.1/142.1 (5hm(dC)), 256.1/140.1 (5f(dC)), 272.1/156.1 (5ca(dC)), 252.1/136.1 (dA), 228.1/112.1 (dC), 268.1/152.1 (dG); 243.1/127.1 (dT); 257.1/136.1 (^13^C_5_-dA I.S.), 246.1/130.1 (13C,^15^N_2_-dT I.S.), 245.1/129.1 (d_3_-5m(dC) I.S.), 261.1/145.1 (d_3_-5hm(dC), I.S.), and 264.1/112.1 (gemcitabine, a dC analog coeluting with 5ca(dC) and used as I.S. for 5f(dC) and 5ca(dC)).

### Immunofluorescence (IF) analysis

#### NPCs

Differentiated neurospheres were washed with PBS, fixed with 4% paraformaldehyde (PFA) for 15 min and permeabilized for 45 min with 0.1% Triton X-100 in PBS. Prior to incubation with the primary antibodies, cells were blocked for 45 min with PBS containing 0.1% Triton X-100, 5% normal goat serum (NGS) (Jackson Immuno Research, 005-000-121), 5% bovine serum albumin (BSA) (Sigma, A2153). Cells were next incubated overnight at 4 °C with primary antibodies (Supplementary Table 3) diluted in 0.1 % Tween-20 + 0.5 % BSA + 0.5 % goat serum in PBS. Prior to the incubation with specific secondary antibodies (Supplementary Table 3) cells were washed 3 x 5 min with PBS containing 0.1% Tween 20. Nuclei were counterstained with DAPI for 10 min. Cells were then visualized using the EVOS FL Auto with 10x objective and images acquired using the TagBFP Light Cube (ThermoFisher Scientific, AMEP4668), GFP Light Cube (ThermoFisher Scientific, AMEP4651) and TexasRed Light Cube (ThermoFisher Scientific, AMEP4655). Images of differentiated NPCs were processed using Fiji software v. 1.54f (NIH). The fluorescent signal from TUJ1 and GFAP was quantified using iMaris software (Oxford Instruments, version 8.2.1). The numbers of voxels were normalized by the number of DAPI positive cells. Per clone, approximately 1000 cells per well, in total 2000 per experiment, were analyzed. Data from four independent experiments are included.

#### Brain Sections

Brain isolations and staining were performed as previously described ^29^. Images of stained sections were captured using the Zeiss AiryScan confocal microscope (Zen Black v. 14.0.12.201, Zeiss) with 40X water immersion objective and analysed using Fiji ImageJ (v.1.53f51, National Institutes of Health). Raw LSM files were imported and stitched using the Grid/Collection Stitching and flattened to 2D image using summed intensity Z projection. The region of interest was outlined using the freehand tool and the raw integrated density measurements of MAP2 and GFAP normalized to the raw integrated density of DAPI. Data is from 1-2 hemispheres of 4 WT and 4 *Aag^-/-^* mice.

#### iNs

Coverslip-seeded iNs were fixed with 4% PFA (1040051000, Merck) for 15 min at RT and washed 3 times for 5min while shaking. The cells were permeabilized with 0.1% Triton- X (T8787 250 ml, Sigma Aldrich) in PBS at room temperature (RT) for 15 min while shaking. Blocking solution containing 5 % BSA (A2153-100G, Sigma Aldrich), 5 % goat serum (005- 000-121, Jackson Immuno Research), 0.1 % Triton-X was prepared in PBS and the cells were blocked for 1 hr with gentle shaking at RT. Primary antibody dilutions (Supplementary Table 3) were prepared in 0.5% BSA, 0.5% goat gut serum, 0.1% Triton-X in PBS and incubated at 4℃ overnight. The next day, coverslips were washed with PBS-T 3 times, 5 min each. 75 µl of secondary antibody dilutions (Supplementary Table 3) in PBS-T were added to the cells and incubated for 60 min at RT. After incubation, the cells were washed with PBS-T 3 times, 5 min each. DAPI solution was added to the wells and incubated for 10 min with shaking. The coverslips were washed with PBS 2 times, 5 min each and finally coverslips were mounted with ProLong™ Gold Antifade Mountant with DNA Stain DAPI (Invitrogen, P36935). The images were taken with Leica SP8 STED Confocal microscope with 20X air objective.

All images were analysed using Fiji ImageJ (v. 1.54f, NIH). For MAP2 and TUJ1 analyses, the summed intensity projection was created. The integrated density of signals was measured and normalized to the number of cells in the image. To analyse the DSB status of cells, DAPI stained nuclei were automatically outlined and saved to the ROI manager using a customized macro. yH2AX foci were selected using Multipoint tool before a mask was created of the points, resulting in an 8-bit black image with single white pixels where the foci were selected. All cell outlined in ROI manager were measured, and the output measurement raw integrated density was divided by 255 to obtain the number of foci counted within each cell.

### Whole Cell Extracts and Immunoblot analysis

The whole cell extracts (WCE) were prepared as described in^26^. Briefly, cell pellets were resuspended in 2 PCV hypotonic lysis buffer (10% glycerol, 20 mM Hepes (pH 7.9), 0.2 mM EGTA, 2 mM MgCl2, 2 mM DTT, 0.1 mM PMSF, 1x Protease Inhibitor Cocktail) and incubated on ice 5 min. Three freeze and thaw cycles were performed and subsequently NaCl and NP- 40 were added to final concentrations 0.5 M and 0.5% (v/v), respectively followed by 20 min incubation on ice. The samples were diluted by addition of 8 PCV hypotonic lysis buffer containing 50 mM NaCl and subjected to sonication using Bioruptor (Diagenode) (5 pulses, 30 sec ON, 30 sec OFF). Samples were centrifuged 10 min, 12,000 x g at 4 °C and the WCE supernatant was collected. For the immunoblot analysis, equal amounts of WCE were ran on NuPage 4-12% Bis-Tris Protein Gel (NP0321, Invitrogen) and proteins transferred onto 0.45 µm nitrocellulose membrane. Upon blocking with Intercept Blocking Buffer (927-70001, Licor) at RT, 1 h, proteins were detected with specific primary antibodies: PAR (Enzo ALX-804-220- R100, 1:1,000), AAG (LSBio-C133325 clone 1E10, 1:500), Tubulin (Cell Signaling, 2144, 1:10,000) and subsequently the corresponding infrared (IR) Dye-conjugated secondary antibodies (Li-COR Biosciences, 827-11081 and 925-32210, 1:15,000). The signal was visualized using direct IR fluorescence with the Odyssey Scanner, LI-COR Biosciences.

### Single cell sequencing (scRNA-seq)

iNs were collected with using papain (Worthington, LK003178) dissolved in 1:1 accutase : DPBS, neutralized with DMEM/F12 with ROCKi and DNase, and pelleted by centrifugation. The pellets were resuspended in DPBS containing 400 µg/ml BSA and cells strained through 35 µm cell strainer. Equal numbers of cells were mixed and subjected to single cell RNA (scRNA) sequencing. scRNA capture and library preparation were performed from four cell suspensions, using the Chromium Next GEM Single Cell 3’ cDNA Kit v3 (10X Genomics, CA; USA), according to the manufacturer’s protocol. In brief, 14,000-20,000 cells were loaded onto the 10X Chromium chip using the recommended loading volumes and run on the 10X Chromium V3 controller (10X Genomics, CA; USA). Following GEM generation on the chip, the Gel Beads were dissolved, primers released, and cells lysed, enabling production of barcoded, full length cDNAs through a cDNA synthesis step. Next, the GEMs were broken, and pooled fractions recovered. Barcoded, full-length cDNAs were amplified via 12 PCR cycles, followed by an AMPure XP beads (Beckman Coulter, Inc., IN, USA) clean up, enzymatic fragmentation and cDNA amplicon size selection. Illumina P5, P7, i7 sample index, and TruSeq read2 (read2 primer sequence) were added via end repair, A-tailing, Adaptor Ligation, and PCR. The final libraries were purified using the AMPure XP (Beckman Coulter, Inc., Indianapolis, IN, USA), quantitated by Collibri Library Quantification Kit (Thermo Fisher Scientific, Waltham, MA, USA) and validated using Agilent DNA 1000 Kit on a Bioanalyzer (Agilent Technologies, Santa Clara, CA, USA). Libraries were normalized and pooled to 350 pM and subjected to clustering on a NovaSeq6000 SP flowcell (Illumina, San Diego, CA, USA). Finally, sequencing was performed for 138 cycles (using the following read geometry: 28 bp for read 1 (cell barcode and UMI), 8 bp for read 2 (sample index) and 101 bp for read 3 (transcript) on a NovaSeq6000 instrument (Illumina, Inc. San Diego, CA, USA), according to the manufacturer’s instructions. Base calling was done on the NovaSeq6000 instrument by RTA 2.4.6. FASTQ files were generated using bcl2fastq2 Conversion Software v2.20.0.422 (Illumina, Inc. San Diego, CA, USA).

### scRNA-seq data analysis

#### Quantification and preprocessing of raw sequencing data

Raw sequencing reads were generated using CellRanger version 7.0.1 and processed with the alevin-fry pipeline. Transcriptomic references were constructed using the splici strategy to incorporate both spliced and unspliced sequences. Reads were aligned and quantified using the alevin module in Salmon version 1.7.0, and molecule information was stored in RAD format. Permit lists for cell barcodes were generated using both filtered and unfiltered strategies, followed by collation of RAD files. Quantification was carried out using the cr-like expectation-maximization resolution mode. Cell calling was further refined with CellBender version 0.2.2 to remove ambient RNA contamination and identify high-confidence cells. Spliced, unspliced, and total gene count matrices were stored in h5ad format and served as input for all downstream analyses.

#### Data integration and preprocessing

Data integration and preprocessing were performed in Python using Scanpy and Scrublet. Prior to integration, low-quality cells and potential doublets were removed. Specifically, cells identified by Scrublet as doublets were excluded, along with cells containing more than 15% mitochondrial reads, fewer than 2,000 or more than 25,000 UMIs, or fewer than 300 detected genes. Genes expressed in fewer than 30 cells were also filtered out. To improve the robustness of pseudotime inference, our single-cell dataset was combined with a publicly available time course dataset of iPSC-derived neurons (ArrayExpress Accession: E-MTAB-10632), using only samples corresponding to day 14 or earlier to match our experimental timespan (days 4–14). Highly variable genes were identified on the combined dataset by selecting the top 2,000 most variable genes using Scanpy.

#### Barcode correction

Batch correction was performed using the scvi-tools package. The combined dataset was corrected for batch effects using scVI, a deep generative model for single-cell data. The model was configured with a negative binomial gene likelihood, a 20- dimensional latent space, one hidden layer, and 64 hidden units, and was trained for 100 epochs. The resulting batch-corrected latent space was used to construct a nearest-neighbor graph to support downstream trajectory inference and visualization.

#### RNA Velocity analysis and Pseudotime inference

Pseudotime inference and RNA velocity analysis were conducted using the dynamo-release package (version 1.2.0). RNA velocity vectors and pseudotime trajectories were inferred using gene-wise kinetic modeling, following the Monocle recipe for gene filtering. A smoothed RNA velocity vector field was used to estimate pseudotime via Hodge decomposition, which captures the global potential landscape of cell state transitions. These analyses were based on the combined dataset, leveraging both our data and the external time course dataset to improve trajectory resolution. Following pseudotime estimation, downstream analyses were restricted to cells originating from our dataset.

#### Differential expression along pseudotime

Differential expression analysis was performed in R using the tradeSeq package. Generalized additive models were fitted to gene expression profiles using five knots, and significance was assessed using the patternTest function. Genes with a patternTest p-value below 0.1 were considered significant. A total of 1,379 genes met this threshold and were included in downstream clustering.

#### Gene module clustering

Gene clustering was performed in R using the clusterExperiment package. Genes with significant pseudotime-dependent expression were grouped into co- expression modules using the RSEC algorithm, evaluated across 30 pseudotime points. Clusters were merged based on adjusted p-values, resulting in 10 distinct gene modules. These modules were used for biological interpretation and gene ontology enrichment analysis.

### mRNA sequencing (RNA-seq)

Total RNA was isolated with RNeasy kit (Qiagen,74106) and RNA sequencing libraries prepared using the Illumina Stranded mRNA prep (ligation) kit (Illumina, San Diego, CA, USA) according to the manufacturer’s protocol. In brief, mRNA was purified from 225 ng total RNA with poly-T oligo-attached magnetic beads, followed by random fragmentation at 94°C for 8 min. cDNAs were first synthesized (42°C for 15 min) using random hexamer oligonucleotides and including actinomycin D. Second strand cDNA synthesis (16°C, 1 h) was performed next using deoxyuridine triphosphate (dUTP) instead of deoxythymidine triphosphate (dTTP). 3’ end adenylation was performed (37°C, 30 min) and anchors ligated to facilitate a second ligation of Illumina dual index adapter oligonucleotides (30°C, 10 minutes). After a fragment cleanup step with AMPure XP (Beckman Coulter, Inc., Indianapolis, IN, USA), library fragments were enriched by 13 cycles of PCR reaction. The final libraries were purified using the AMPure XP (Beckman Coulter, Inc., Indianapolis, IN, USA), quantitated by KAPA Library Quantification Kit (Kapa Biosystems, Inc., Wilmington, MA, USA) and validated by Agilent High Sensitivity DNA Kit on a Bioanalyzer (Agilent Technologies, Santa Clara, CA, USA). DNA fragments were in the range of 200-600 bp and peaked around 301 bp. Libraries were pooled and normalized to 1.5 pM and subjected to clustering on a NovaSeq6000 SP flowcell (Illumina, San Diego, CA, USA). Finally, paired end read sequencing was performed for total 138 cycles (2X59 bp for the insert and 2X10 bp for the indexes) on a NovaSeq6000 instrument (Illumina, Inc. San Diego, CA, USA). Base calling was done on the NovaSeq6000 instrument by RTA 2.4.6. FASTQ files were generated using bcl2fastq2 Conversion Software v2.20.0.422 (Illumina, Inc. San Diego, CA, USA).

### RNA-seq data analysis

#### Data acquisition and processing

Raw FASTQ files were assessed for quality by using FastQC (v0.11.9, https://www.bioinformatics.babraham.ac.uk/projects/fastqc/). Reads were filtered and trimmed with fastp (v0.20.1, https://github.com/OpenGene/fastp) and sequences aligned against the human reference genome (GRCh38) by using STAR^57^ (v2.7.9a, https://github.com/alexdobin/STAR). Post-alignment quality metrics were extracted with Picard CollectRNASeqMetrics (v2.21.5). Transcript-level abundance estimates were generated via Salmon^58^ (v1.7.0, https://combine-lab.github.io/salmon/). The resulting transcript counts were aggregated to gene-level counts in R, by using tximport (v1.20.0). Publicly available RNA-seq data from Bioprojects PRJNA704023^10^ and PRJNA865253^9^ were obtained from the Sequence Read Archive (SRA) and processed as described above, to ensure comparability with our data.

#### Data Normalization and Batch Correction

Gene-level counts were initially filtered with HTSfilter (v1.42.0) in the R statistical environment. Batch effects between datasets were corrected with ComBat from the sva package (v3.50.0).

#### Differential Expression Analysis

Differential gene expression analysis was performed with DESeq2^59^ (https://bioconductor.org/packages/DESeq2/). Significance was determined by comparing the genotype-specific differences at each differentiation time point against the baseline genotype difference at day 0, with genes considered significantly differentially expressed based on an adjusted p-value < 0.05 and an absolute log fold-change (logFC) > 0.1. DAVID v6.8 gene-ontology (GO) enrichment analysis was performed to identify enriched Biological Processes (BP) across gene clusters.

#### Heatmap Visualization

The log fold-change values of significant differentially expressed genes were visualized by the ComplexHeatmap package (v2.18.0) in R. Default settings were applied, and genes were clustered into 5 distinct groups using consensus k-means clustering with 1500 iterations.

### Proximity Ligation Assay (PLA)

The stem cells grown on coverslips were fixed with 4 % PFA for 15 min at RT, on D0 and D7 of differentiation protocol. The coverslips were next washed 3 times with PBS and permeabilised with 0.1 % Triton-X in PBS for 15 min. Duolink proximity ligation assay (Sigma Aldrich, DUO92101) was performed according to manufacturer’s protocol with AAG (1:80, Mouse, LSBio-C133325 clone 1E10) and TDG (1:50, Polyclonal rabbit, Invitrogen PA5-29140) primary antibodies. The D0 images were taken with Evos FL Auto 2 with z-stack size of 1 µm by using 40x objective. The D7 images were taken with Zeiss AiryScan confocal microscope using 40x objective (Zen Black v. 14.0.12.201, Zeiss).

### DNA Glycosylase Assay

Recombinant full-length human AAG and mouse TDG were obtained by: expression from pET19b-PPS-AAG and pET28-mTDG plasmids in *E. coli* BL21(DE3) cells, respectively, and purification according to the established procedure^60, 61^. For the denaturing gel analysis of DNA glycosylase products, the reactions (10μl) containing: 50 mM Tris-HCl at pH8, 1 mM DTT, 1 mM EDTA and 0.1 mg/ml BSA and 20 nM FAM-labelled 5fC hairpin DNA substrate (Supplementary Table 1) were incubated with varied amounts (0.05, 0.1, 0.2, 0.4 or 0.8 μM) of AAG or recombinant BSA for 5 min on ice, followed by addition of 0.2 μM mTDG and incubation 1 h at 37 °C. Reactions were quenched with 0.2N NaOH and heated 15 min at 75 °C. Formamide loading buffer was added to each sample and cleaved products resolved on a 12% denaturing polyacrylamide gel. Products were visualized by ChemiDoc Imaging System (Bio-Rad) and analysed by Fiji software v. 1.54f (National Institutes of Health).

### Single Molecule Analysis

Single-molecule analysis of DNA-binding proteins from nuclear extracts (SMADNE) was performed as described previously^38, 62, 63^, with few modifications. To generate DNA template containing a single 5fC modification in the middle of 12.6kb template, DNA tethering kit (Lumicks) was used by ligating a double stranded DNA oligo with 5fC (sequences listed in the Supplementary Table 1) into DNA arms as described in^64^. Nuclear extracts expressing fluorescently tagged AAG or TDG, were obtained by transient CaCl_2_ transfection of HEK293T cells with pEGFP.N1-AAG^26^ and pHTN TDG-Halo CMV-neo plasmids, respectively. 48hr after transfection, cells expressing TDG-HaloTag were labelled with saturating JF-635 HaloTag ligand, then collected, and nuclear extracts prepared with the nuclear extraction kit (Abcam, ab113474) according to manufacturer’s protocol, with inclusion of 3 x 10 sec sonication. Total protein concentration was determined with Bradford assay (Bio-Rad), while <u>Quant-</u> iT™PicoGreen™ dsDNA Assay Kit (Invitrogen, P11496) was used to determine the nucleic acid concentration (average dsDNA concentration was ∼200 ng/ul and total protein ∼2 mg/ml). The efficiency of target protein overexpression was confirmed by immunoblot analysis (Supplementary Fig. 5a-b), following procedure described above and in^38^. The single-use aliquots were flash-frozen in liquid nitrogen and kept at -80 °C.

Single molecule analysis was performed with C-Trap instrument (Lumicks). Prior to each experiment, the flow cell was cleaned and passivated with 2% BSA. Experiment reagents were diluted in the recording buffer (25 mM HEPES, pH 7.5, 125 mM NaCl, 0.1 mg/ml BSA, 1 mM DTT, and 1 mM Trolox). Channels were loaded with: 1) streptavidin-coated beads (Ø 1.76 μm, 1:2000); 2) biotinylated 12.6kb DNA containing a single 5fC (1:400); 3) recording buffer; and 4) nuclear extract (diluted 1:20-1:50 for TDG-HaloTag and 1:225-1:350 for AAG-EGFP). A single DNA tether was formed between optically trapped beads using 15% overall laser power by adjusting the distance between the traps while monitoring the force-distance curve. The tether was subsequently washed in the recording buffer, after which the flow was stopped and the DNA stretched to 10 pN. Kymograph recordings were then acquired for 5-15 min as the tether was moved into the extract channel. Imaging was performed using continuous line scanning with a pixel size of 100 nm, a pixel dwell time of 0.1 ms, and a line time of 30 ms. AAG-EGFP and TDG-HaloTag JF-635 were excited using 488 nm and 638 nm lasers at 5% and 4.9% power, respectively. Emission was collected using 525/45 nm and 680/42 nm bandpass filters with a 60× NA 1.2 water immersion objective, and photons detected using single-photon avalanche photodiodes.

Kymographs were analyzed using the Python package Pylake (v1.5.3, LUMICKS), following procedures described in^38^. Briefly, line tracking of binding events was performed using Gaussian fit to determine the positions. Binding events occurring within ±0.1 μm of the DNA midpoint (the 5fC site) were classified as specific binding events (Fig. 5). The time between two consecutive specific binding events was defined as the gap time. Binding and gap times were analyzed using cumulative residence time distributions (CRTD) and cumulative gap time distributions (CGTD). The data were fit using either a one-phase exponential decay or in the case of TDG-specific binding a two-phase exponential decay model, as described in, to obtain rate constants. Photobleaching measurements were performed and analyzed as described in^38^, and the resulting photobleaching rates applied to correct the dwell times. Protein concentrations in each kymograph were determined by measuring the background fluorescence intensity and extrapolating to a standard curve of known fluorophore concentrations, which was used to determine the association rate. To calculate the apparent association rate to the 5fC sites (*k_on,app_*) (Fig. 5 and Table 1) CGTD measurements normalized to protein concentration were used. The dissociation constant (*K_d_*) was calculated from the 5fC specific dissociation rate (*k_off_* = 1/τ) and the association rate, according to equation *K_d_* = *k_off_*/*k_on_*.

## Statistical Analysis

Analysis of data was performed using GraphPad Prism v10 (GraphPad Software, Inc., La Jolla, CA). Significances of changes were determined by using Student t test (Figs 1b-i, 4g-h, Supp Fig. 1a-k), Mann-Whitney test (Fig 4d, Supplementary Fig 4b), one-way ANOVA followed by Tukey’s comparison test (Fig 4b) or two-way ANOVA followed by Tukey’s comparison test (Fig 2h-i, Supp Fig. 2c-f, 3b-d, 4d-e), as further indicated in the figure legends. All data represent mean ± SEM; \**p* ≤ 0.05; \*\**p* ≤ 0.01; \*\*\**p* ≤ 0.001; \*\*\*\**p* ≤ 0.0001 vs. control; NS, not significant.

## Data availability

A reporting summary is available as a Supplementary Information file. The scRNA sequencing data reported in this paper are available in GEO under accession GSE326181for iNs and the RNA sequencing data available under accession GSE326180 for stem cells and iNs.

## Acknowledgements

This work has been supported by the Research Council of Norway grant (ID: 335324) to B.v.L. We thank to Keith Caldecott for kindly providing iNs system, as well as for the discussions and feedbacks to the manuscript; to Rossana Aprigliano and Stefano Bradamante for help with the initial NPCs proliferation and differentiation experiments; to Nina-Beate Liabakk for purifying TDG; and to Leona D. Samsons for the discussions. Imaging was performed at the Cellular and Molecular Imaging Core Facility (CMIC), and LC-MS/MS analysis of DNA modification at the Proteomics and Modomics Experimental Core facility (PROMEC), both at NTNU.

## Author Contributions

B.v.L. had the original idea, supervised the project and together with E.M.A, K.G. and A.G. interpreted the experiments. B.v.L., E.M.A and K.G. wrote the manuscript and with A.G. generated figures. E.M.A. performed most of the experiments; K.G. performed brain isolations, imaging, IF analysis, contributed to bioinformatics analysis and together with M.S. and B.V.H. contributed to SMADNE. A.G. performed single molecule measurements and the analysis. D.L.B. performed neurosphere assays. R.D. contributed to *in vitro* DNA glycosylase assays. A.F. together with P.S. performed most of the bioinformatics analysis. N.E.Z. contributed to iN differentiation and RT-qPCR experiments. C.B.V. performed LC-MS/MS analysis of the epigenetic marks.

## Competing interests

The authors declare no competing interests.

## Materials & Correspondence

Correspondence and material requests should be addressed to Barbara van Loon.

## Notes

### Competing Interest Statement

The authors have declared no competing interest.

