## SupplementalInfo for "Alkyladenine DNA glycosylase regulates TDG-mediated demethylation and neuronal gene expression by competitive substrate binding"

**Aksu M.E., et al.**

**SUPPLEMENTARY TABLES AND FIGURES**

**Supplementary Table 1.** List of oligonucleotides used for CRISPRi, DNA glycosylase assays and ligation into the arms for single molecule SMADNE experiments. FAM: 5' 6-FAM (Fluorescein); 5fC: 5-formylcytosine.

| **DNA oligonucleotides** | **Method** | **Sequence 5' to 3'** |
|---|---|---|
| Top non-targeting sgRNA | CRISPRi | ttgGCCCCTGACCGGCACCCTCGgtttaagagc |
| Bottom non-targeting sgRNA | CRISPRi | ttagctcttaaacCGAGGGTGCCGGTCAGGGGCcaacaag |
| Top AAG targeting sgRNA | CRISPRi | ttgGTGGGACTCGGACCCCGGCGgtttaagagc |
| Bottom AAG targeting sgRNA | CRISPRi | ttagctcttaaacCGCCGGGGTCCGAGTCCCACcaacaag |
| 5fC hairpin DNA | DNA glycosylase | [FAM]-CGACGCTATTCA[5fC]GCGTGCTCTGAGGGTTTT CCCTCAGAGCACGCGTGAATAGCGTCG |
| Top strand 5fC | SMADNE | [phosphate]-CAACACCAGTCCATCGCTCA[5fC]GTACAGAGCTG |
| Bottom strand | SMADNE | [phosphate]-ACCACAGCTCTGTACGTGAGCGATGGACTGGT |

**Supplementary Table 2.** List of oligonucleotides used for RT-qPCR analysis of mRNA level.

| **cDNA** | | |
|---|---|---|
| **Name** | **Forward / Reverse** | **Sequence 5' to 3'** |
| ACTB | Fw | GTTACAGGAAGTCCCTTGCCATCC |
| ACTB | Rv | CACCTCCCCTGTGTGGACTTGGG |
| OCT4 | Fw | TCAGGAGATATGCAAAGCAGAAAC |
| OCT4 | Rv | CTGATCTGCTGCAGTGTGGG |
| AAG | Fw | CGGAGTTCTGTGCCATTAGG |
| AAG | Rv | GGTTGGAGTTCTTCGACCAG |
| RBFOX3 | Fw | CCAAGCGGCTACACGTCTC |

| | | |
|---|---|---|
| RBFOX3 | Rv | CGTCCCATTCAGCTTCTCCC |
| SLC17A7 | Fw | CTGGGGCTACATTGTCACTCA |
| SLC17A7 | Rv | GCAAAGCCGAAAACTCTGTTG |
| DCX | Fw | ACGATGAACCCCGACAGA |
| DCX | Rv | GTTGGGATTGACATTCTTGGTG |
| MAP2 | Fw | CAGGAGACAGAGATGAGAATTCC |
| MAP2 | Rv | CAGGAGTGATGGCAGTAGAC |
| PAX6 | Fw | GCCCTCACAAACACCTACAG |
| PAX6 | Rv | TCATAACTCCGCCCATTCAC |
| TDG | Fw | TGACCAAGACTCTCCCCGATA |
| TDG | Rv | CCAGGTCCAGGGTAATGATGC |
| TET1 | Fw | CAGAACCTAAACCACCCGTG |
| TET1 | Rv | TGCTTCGTAGCGCCATTGTAA |
| PARP1 | Fw | CGGAGTCTTCGGATAAGCTCT |
| PARP1 | Rv | TTTCCATCAAACATGGGCGAC |
| RIMS4 | Fw | GGATGTGGAGATCGGTCTGC |
| RIMS4 | Rv | GCGAGCGACTTTGGTCTTCT |
| PDLIM4 | Fw | TGATGACAGCAAGGCTCAGGCA |
| PDLIM4 | Rv | AGGCTTGGTCTGCCATCTTCTG |
| SLC6A3 | Fw | CCACCATGCCATACGTGGTC |
| SLC6A3 | Rv | CCGGTAGAAGTCAACGCTCA |

**Supplementary Table 3.** List of antibodies used in immunofluorescence analysis

| Antibody | Cat# | Dilution |
|---|---|---|
| Anti-GFAP Rabbit Polyclonal Antibody | Aligent Dako Z0334 | NPCs: 1:600<br>Brain: 1:500 |
| Anti-TUJ1 Mouse IgG2a Antibody | Neuromics MO15013 | NPCs: 1:600 |
| Anti-TUJ1 Mouse IgG2a Antibody | Covance MMS-435P | iNs: 1:750 |
| Anti-MAP2 Chicken IgY Antibody | Abcam ab5392 | Brain: 1:500<br>iNs: 1:500 |

| | | |
|---|---|---|
| Anti-phospho-Histone H2A.X (Ser139), JBW301 Mouse IgG1 Antibody | Sigma Aldrich 05-636 | iNs: 1:50 |
| Goat Anti-Mouse IgG1, Alexa Fluor 488 | Invitrogen A21121 | iNs: 1:500 |
| Goat Anti-Mouse IgG2a, AlexaFluor 488 | Invitrogen A21131 | NPCs: 1:500<br>iNs: 1:500 |
| Goat Anti-Rabbit IgG (H+L), Alexa Fluor 594 | Invitrogen A11037 | NPCs: 1:500<br>Brain: 1:500 |
| Goat Anti-Chicken IgY H&L, Alexa Fluor 647 | Abcam ab150175 | iNs: 1:500<br>Brain: 1:500 |
| DAPI solution (1 mg/mL) | ThermoFisher 62248 | 1:1000 |

**Figure S1. Loss of AAG impairs neuronal differentiation in mouse hippocampus, without affecting self renewal and proliferation of neuronal progenitor cells. a** Relative number of WT and Aag knock-out (*Aag*$^{-/-}$) spheres forming neural progenitor cells (NPCs) from two different clones per genotype at day 10. **b** PrestoBlue of WT and *Aag*$^{-/-}$ NPCs proliferation capacity, across five days (n=3, each experiment includes two clones per genotype). **c** Representative immunofluorescence images of Map2 (mature neuronal marker), Gfap (glial marker) and DAPI staining in the postnatal day 5 (P5) WT and *Aag*$^{-/-}$ mouse hippocampus (HPC). Scale bar 100 µm. **d-g** Quantification of Map2 signal intensities from images as the one depicted in (c) across the total HPC (d) and in the specific HPC regions: dentate gyrus (DG) (e), CA3 (f) and CA1 (g). **h-k** Quantification of Gfap signal intensities in the total HPC (h) and in the specific HPC regions: dentate gyrus (DG) (i), CA3 (j) and CA1 (k). (n = 4 animals, 1-2 hemispheres). All error bars represent mean ± SEM , two-tailed Student's t-test, *p ≤ 0.05; **p ≤ 0.01.

**Figure S2. Expression of stemness and neurodevelopmental genes in differentiating induced neurons lacking AAG. a** Relative *AAG* expression levels in non-targeting sgRNA (WT) and AAG targeting sgRNA (sgAAG) stem cells at day 0 (D0) and induced neurons (iNs) at D4, D7 and D14 of differentiation; normalized to β-actin mRNA. **b** Immunoblot analysis of AAG protein levels in D0, D4, D7 and D14 WT and sgAAG iNs, with tubulin as loading control. **c-f** Gene expression levels of *OCT4* (c), *PAX6* (d), *DCX* (e) and *MAP2* (f) in D0, D4, D7 and D14 WT and sgAAG iNs determined by RT-qPCR, n=4**.** All error bars represent mean ± SEM. two-way Anova, Tukey`s multiple comparison test, *p ≤ 0.05; **p ≤ 0.01; ***p ≤ 0.001; ****p ≤ 0.0001.

**Figure S3. AAG and TDG are proximity in iNs, with loss of AAG resulting in increased expression of synaptic genes. a** Relative gene abundance of cluster 1 (from Fig. 3a) marker gene *SLC6A3* during neuronal differentiation in sgAAG vs WT in the left, and sgTDG vs WT in the right panel. **b-d** Gene expression analysis of *SLC17A7* (b), *PDLIM4* (c) and *SLC6A3* (d) in WT and sgAAG stem cells and iNs determined by RT-qPCR, n=4, two-way Anova, Tukey`s multiple comparison test. **e** Proximity Ligation Assay (PLA) showing AAG and TDG in the close proximity, in D0 stem cells (Scale bar 50 µm). All error bars represent mean ± SEM. two-way Anova, Tukey`s multiple comparison test, *p ≤ 0.05; **p ≤ 0.01; ***p ≤ 0.001; ****p ≤ 0.0001.

**Figure S4. AAG lacking iNs have comparable to WT levels of upstream DNA demethylation substrates and TDG expression. a** Representative image of immunoblot analysis targeting poly ADP ribose (PAR) in WT and sgAAG at D0 iNs. Tubulin serves as loading control. **b** Quantification of PAR levels (from n=3 experiments as the one in (a). **c-d**

LC-MS/MS analysis of active DNA demethylation substrates 5-methylcytosine (5mC) (c) and 5-hydroxymethylcytosine (5hmC) (d); n ≥ 3, **e** Gene expression analysis of *TDG* in WT and sgAAG at day 0 (D0) and induced neurons (iNs) at D4, D7 and D14 of differentiation; normalized to β-actin mRNA; n=3. Error bars represent mean ± SEM, one-tailed Student`s t-test followed by Mann-Whitney test (b), two-way Anova, either followed by Bonferroni`s (c-d), or Tukey`s multiple comparison test (e).

**Figure S5. Analysis of protein overexpression in nuclear cell extracts and colocalization of AAG-EGFP and TDG-Halo on 5fC. a** Immunoblot visualization of TDG-Halo nuclear cell extracts (NCE) probed with TDG antibody and Janelia Fluor 635 HaloTag ligand, total protein staining serves as loading control. Purified mouse recombinant TDG (mTDG) is used as loading control. **b** Immunoblot analysis of AAG-EGFP NCE probed with AAG primary antibody, total protein staining used as loading control. Purified human recombinant AAG lacking 80 N-terminal amino acids servs as loading control. **c** Percent of binding events from experiments as the one in Fig. 5e, when AAG alone (green), TDG alone (red) or both AAG and TDG (yellow) reside on 5fC site. **d** Quantification of types of AAG-EGFP and TDG-Halo co-occupancy events in the kymographs as the one presented in Fig. 5e.

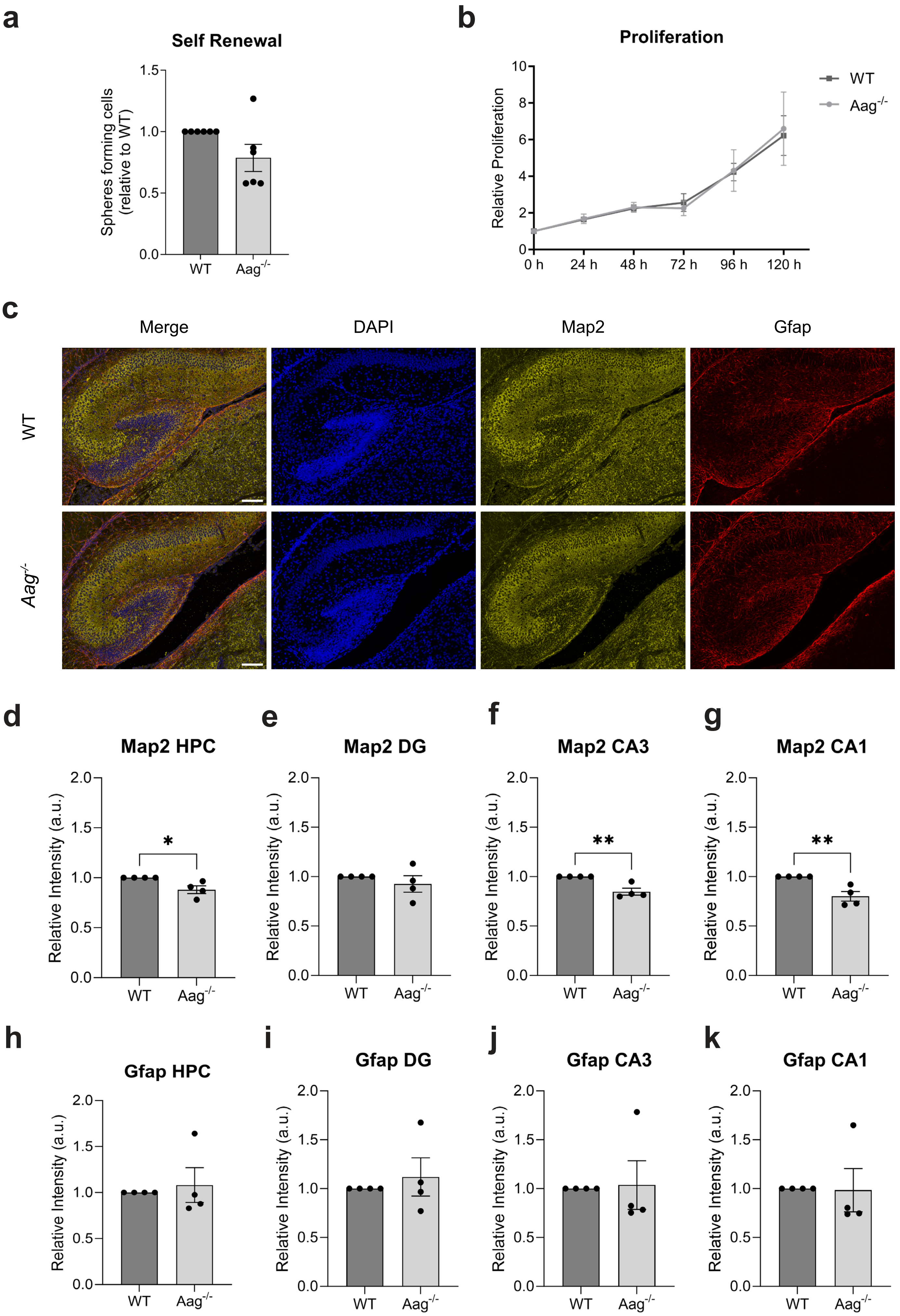
a
Self Renewal
Spheres forming cells (relative to WT)
1.5
1.0
0.5
0.0
WT
Aag-/-
b
Proliferation
Relative Proliferation
10
8
6
4
2
0
0 h
24 h
48 h
72 h
96 h
120 h
WT
Aag-/-
c
Merge
DAPI
Map2
Gfap
WT
Aag-/-
d
Map2 HPC
e
Map2 DG
f
Map2 CA3
g
Map2 CA1
Relative Intensity (a.u.)
2.0
1.5
1.0
0.5
0.0
*
**
WT
Aag-/-
h
Gfap HPC
i
Gfap DG
j
Gfap CA3
k
Gfap CA1

Figure S1.

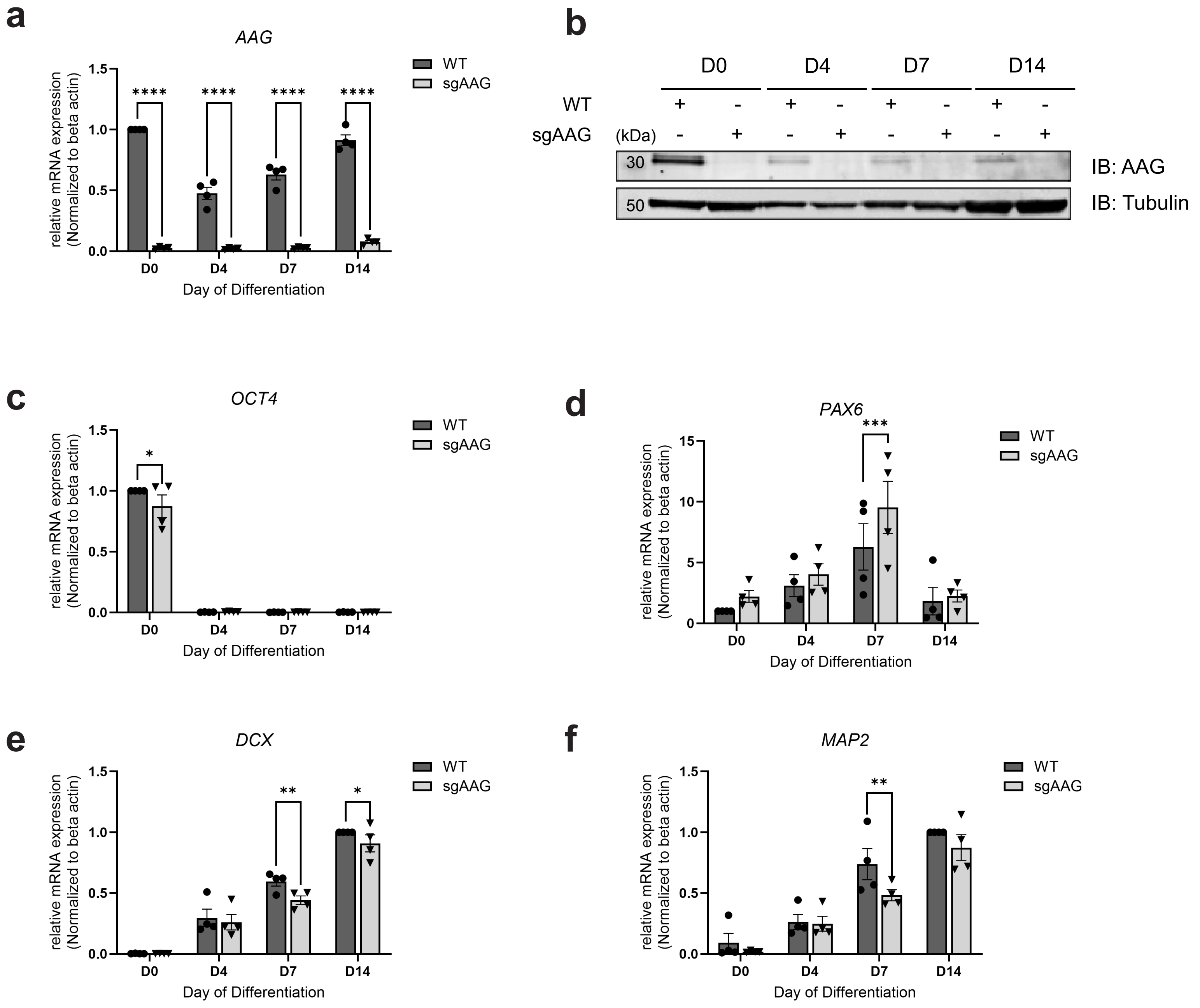
a
AAG
relative mRNA expression (Normalized to beta actin)
Day of Differentiation
WT
sgAAG
b
D0
D4
D7
D14
(kDa)
IB: AAG
IB: Tubulin
c
OCT4
d
PAX6
e
DCX
f
MAP2

Figure S2.

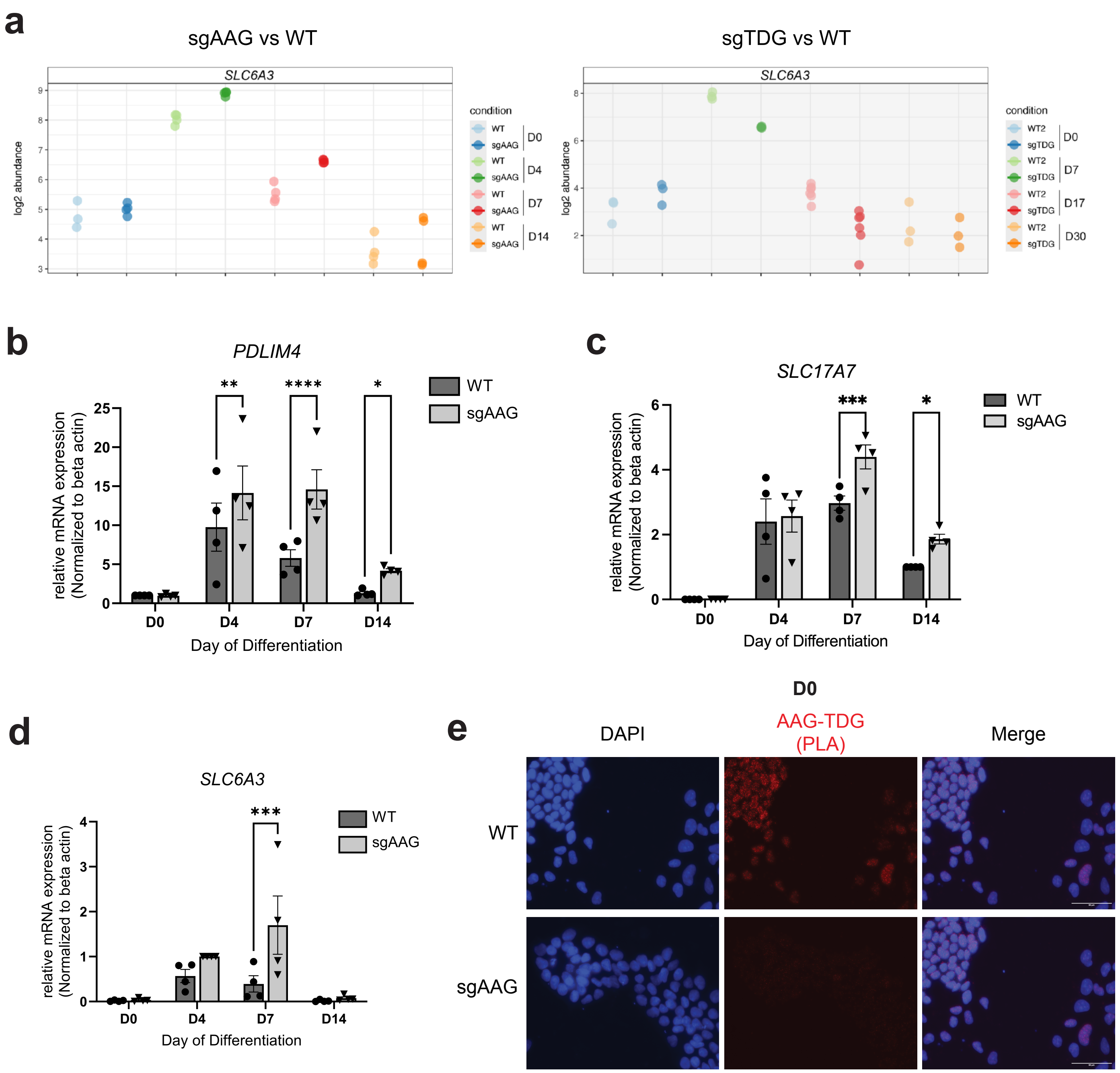
a
sgAAG vs WT
SLC6A3
log2 abundance
condition
WT
sgAAG
D0
D4
D7
D14
sgTDG vs WT
WT2
sgTDG
D17
D30
b
PDLIM4
relative mRNA expression
(Normalized to beta actin)
Day of Differentiation
c
SLC17A7
d
SLC6A3
e
DAPI
AAG-TDG
(PLA)
Merge

**Figure S3.**

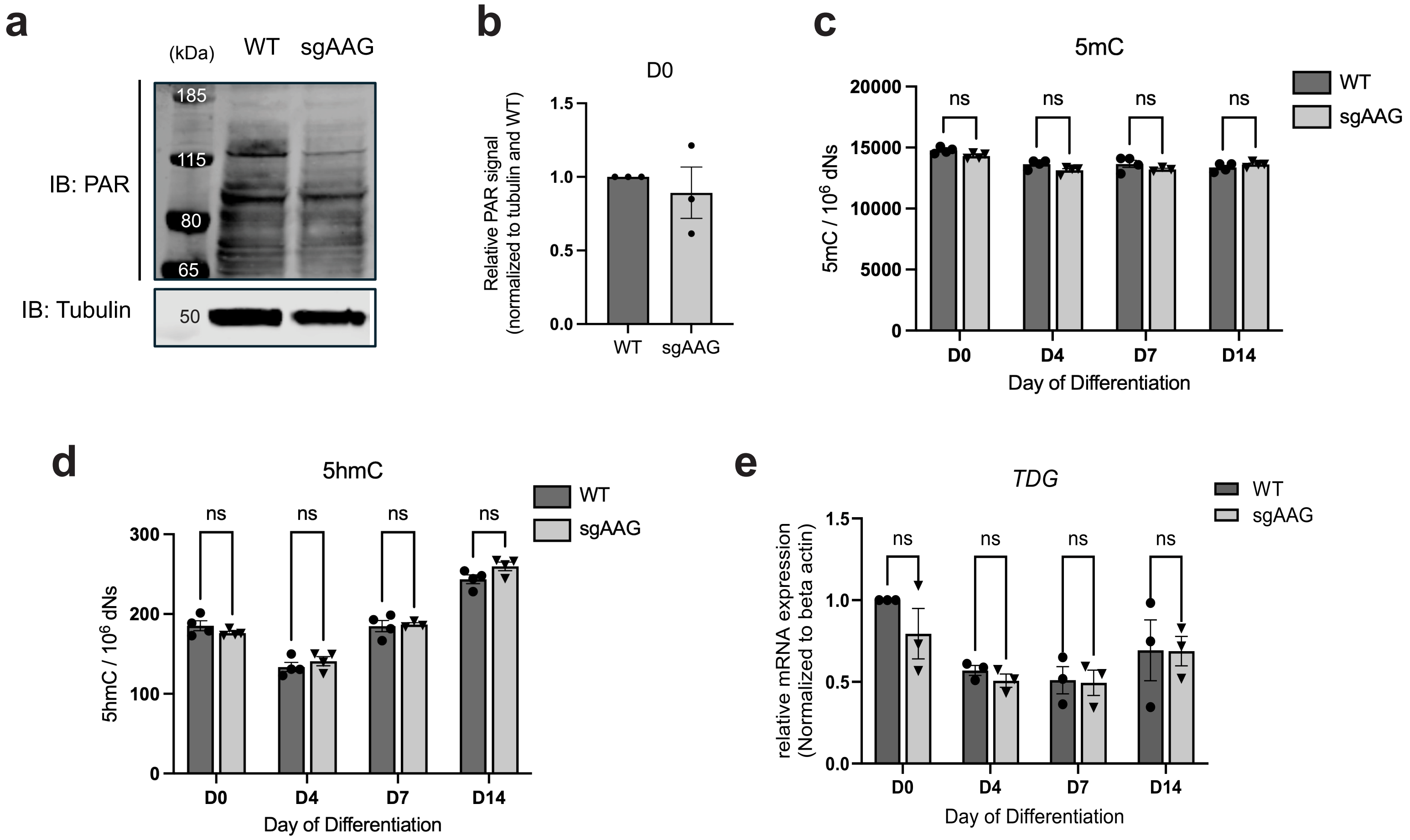
a
(kDa)
WT
sgAAG
IB: PAR
185
115
80
65
IB: Tubulin
50
b
D0
Relative PAR signal (normalized to tubulin and WT)
1.5
1.0
0.5
0.0
WT
sgAAG
c
5mC
WT
sgAAG
20000
15000
10000
5000
0
5mC / 10^6 dNs
ns
D0
D4
D7
D14
Day of Differentiation
d
5hmC
WT
sgAAG
300
200
100
0
5hmC / 10^6 dNs
ns
D0
D4
D7
D14
Day of Differentiation
e
TDG
WT
sgAAG
relative mRNA expression (Normalized to beta actin)
1.5
1.0
0.5
0.0
ns
D0
D4
D7
D14
Day of Differentiation

Figure S4.

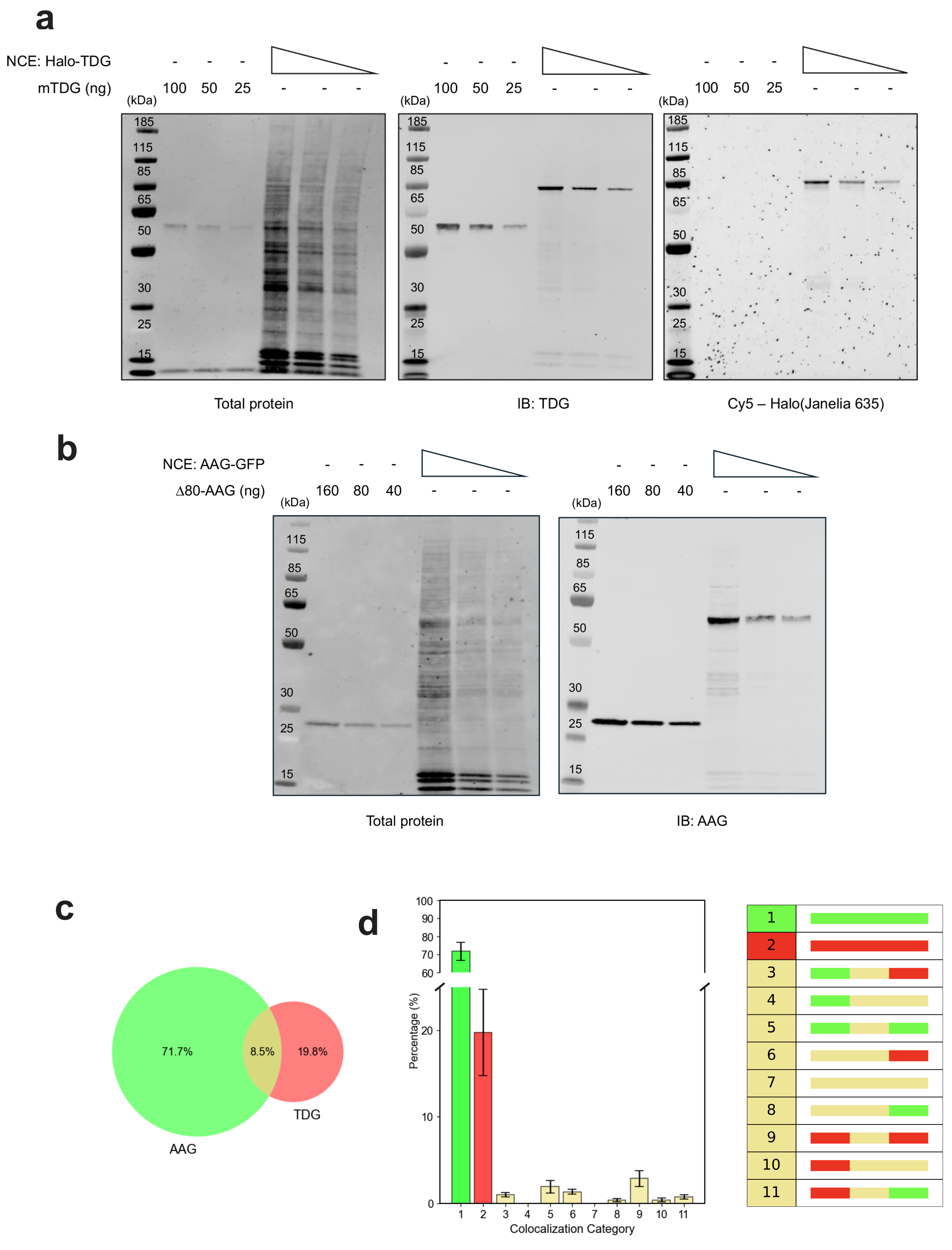

a
NCE: Halo-TDG
mTDG (ng)
100 50 25
(kDa)
185
115
85
65
50
30
25
15
Total protein
IB: TDG
Cy5 – Halo(Janelia 635)
b
NCE: AAG-GFP
Δ80-AAG (ng)
160 80 40
Total protein
IB: AAG
c
71.7%
8.5%
19.8%
AAG
TDG
d
Percentage (%)
Colocalization Category
1 2 3 4 5 6 7 8 9 10 11


Figure S5.
